# Strong and specific connections between retinal axon mosaics and midbrain neurons revealed by large scale paired recordings

**DOI:** 10.1101/2021.09.09.459396

**Authors:** Jérémie Sibille, Carolin Gehr, Jonathan I. Benichov, Hymavathy Balasubramanian, Kai Lun Teh, Tatiana Lupashina, Daniela Vallentin, Jens Kremkow

## Abstract

The superior colliculus (SC) is a midbrain structure that plays important roles in visually guided behaviors. Neurons in the SC receive afferent inputs from retinal ganglion cells (RGC), the output cells of the retina, but how SC neurons integrate RGC activity *in vivo* is unknown. SC neurons might be driven by strong but sparse retinal inputs, thereby reliably transmitting specific retinal functional channels. Alternatively, SC neurons could sum numerous but weak inputs, thereby extracting new features by combining a diversity of retinal signals. Here, we discovered that high-density electrodes simultaneously capture the activity and the location of large populations of retinal axons and their postsynaptic SC target neurons, permitting us to investigate the retinocollicular circuit on a structural and functional level *in vivo*. We show that RGC axons in the mouse are organized in mosaics that provide a single cell precise representation of the retina as input to SC. This isomorphic mapping between retina and SC builds the scaffold for highly specific wiring in the retinocollicular circuit which we show is characterized by strong connections and limited functional convergence, established in log-normally distributed connection strength. Because our novel method of large-scale paired recordings is broadly applicable for investigating functional connectivity across brain regions, we were also able to identify retinal inputs to the avian optic tectum of the zebra finch. We found common wiring rules in mammals and birds that provide a precise and reliable representation of the visual world encoded in RGCs to neurons in retinorecipient areas.

**HIGHLIGHTS:**

- High-density electrodes capture the activity of afferent axons and target neurons *in vivo*
- Retinal ganglion cells axons are organized in mosaics
- Single cell precise isomorphism between dendritic and axonal RGC mosaics
- Midbrain neurons are driven by sparse but strong retinal inputs
- Functional wiring of the retinotectal circuit is similar in mammals and birds

## INTRODUCTION

Retinal ganglion cells (RGCs) encode the visual world in over 30 parallel functional pathways (Baden et al., 2016) and send this information via axons along the optic nerve to multiple and distributed areas in the vertebrate brain (Figure 1A) (Martersteck et al., 2017). A major retinorecipient area in rodents is the superior colliculus (SC) in the midbrain (Ellis et al., 2016; Kremkow and Alonso, 2018), referred to as optic tectum (OT) in non-mammalian vertebrates. The SC is an evolutionary old brain structure that is part of the extrageniculate visual pathway (Beltramo and Scanziani, 2019) and central for visually guided behaviors (Basso et al., 2021; Isa et al., 2021). While we have learned a lot about how SC neurons process visual stimuli (reviewed in (Cang et al., 2018)), how SC neurons integrate retinal activity on a functional level *in vivo* is still largely unknown.

**Figure 1:**
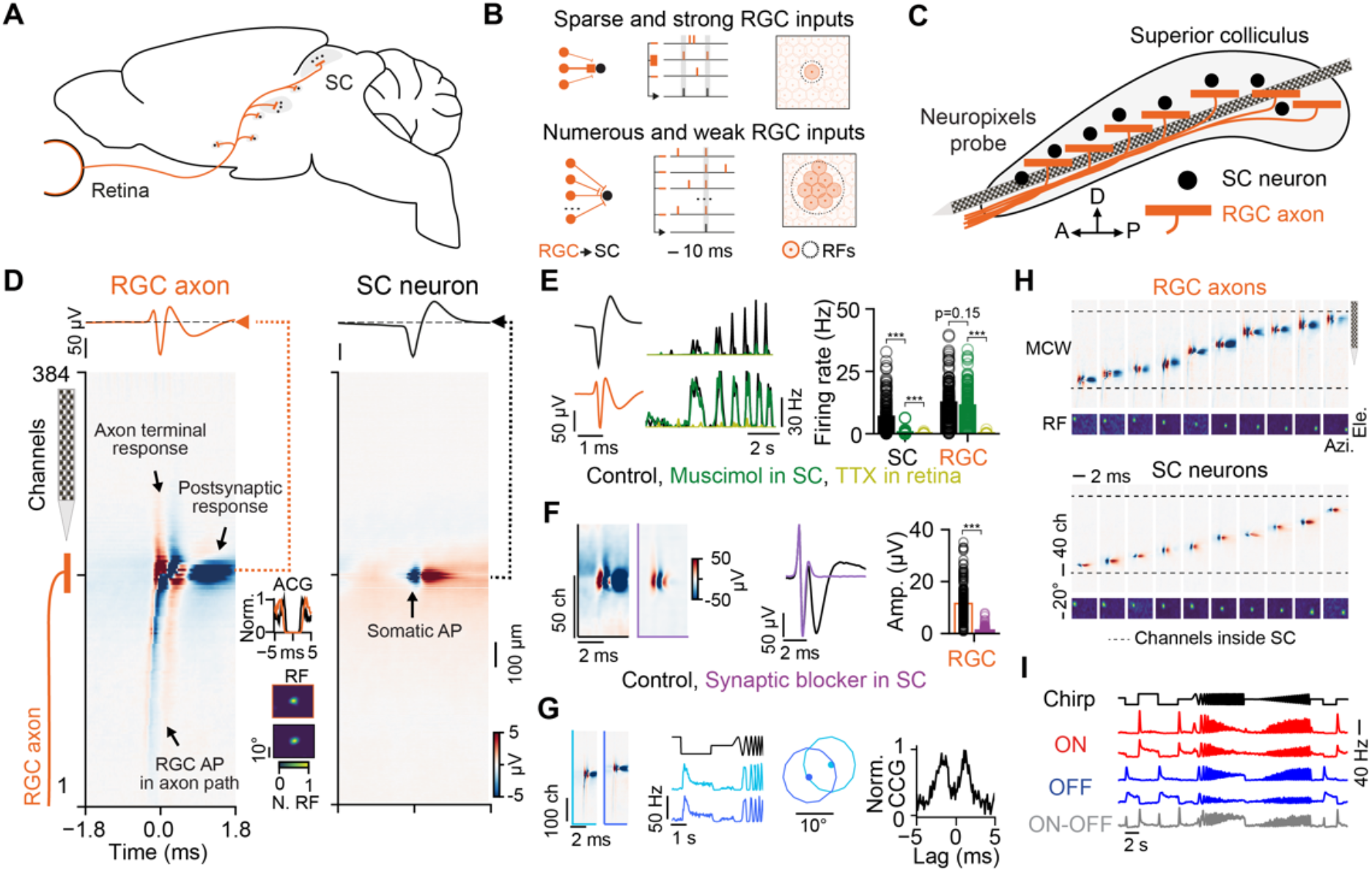
Simultaneous recordings of RGC axons and local neurons in the mouse SC. (A) Retinofugal projections in the mammalian brain. The superior colliculus (SC) is a major brain structure of the visual system receiving afferent inputs from retinal ganglion cells (RGC), the output cells of the retina. (B) Possible mechanisms of how SC neurons integrate RGC inputs *in vivo*. SC neurons could be driven by sparse but strong RGC inputs (top) or by the synchronous activation of numerous weak RGC inputs (bottom). The schematic shows the possible retinocollicular wiring (left), synaptic integration (middle) and connection specificity (right). (C) Experimental setup for simultaneous recording of RGC axons and SC neurons with high-density electrodes (Neuropixels probes) in the mouse SC *in vivo*. (D) Average spatiotemporal electrical signal of an RGC axonal action potential (AP) (left, RGC axon) and somatic SC AP (right, SC neuron) along the Neuropixels probe. The RGC AP propagating along the path of the RGC axon is visible in the multi-channel waveforms (MCW) together with the axon terminal response and the evoked postsynaptic response. ACG = spike train auto-correlogram; RF = receptive field. (E) Pharmacological confirmation of axonal and somatic waveforms. Left, visually evoked activity of an example SC neuron (top) and RGC axon (bottom) during different pharmacological conditions shown in the peri-stimulus time histograms (PSTH): control condition (black), after local Muscimol application in the SC (green) and after injecting tetrodotoxin (TTX) in the eye (light green). Right, Average firing rates during the different conditions (n = 5 mice, n = 224 SC neurons, n = 215 RGC axons). (F) Pharmacological confirmation the second trough in RGC axonal waveforms is postsynaptic evoked activity (n = 3 mice, n = 203 RGC axons). (G) Recording from neighboring RGCs of the same functional type and RGC mosaic, identified by similarity in visually evoked responses to a chirp stimulus (PSTH), non-overlapping RF centers and putative electrical coupling, evident by the double peaks in the spike train cross-correlogram (CCG). (H) Simultaneously recorded RGC axons and SC neurons cover a large part of the SC circuit and different retinotopic positions. The dashed black lines enclose the recording sites located within the SC. (I) Functional diversity of the recorded RGC axons identified by responses to a chirp stimulus. Shown are PSTHs of five distinct functional RGC types: ON-transient, ON-sustained, OFF-transient, OFF-sustained and ON-OFF. For all panels: *** = p < 0.001.

There are multiple possible mechanisms how SC neurons could integrate RGC inputs. SC neurons might be driven by sparse but strong RGC inputs (Figure 1B, top) such that individual RGCs can drive SC spiking. Alternatively, SC neurons could receive numerous but weak RGC inputs and simultaneous activation of multiple pre-synaptic RGCs is required to drive SC spiking (Fig. 1B, bottom). These two distinct wiring schemes have implications for how SC neurons represent the visual world encoded in the diverse pathways of their retinal afferents. Strong but sparse inputs would indicate that SC neurons reliably represent the activity of specific retinal pathways, comparable to the retinogeniculate circuit (Rosón et al., 2019; Usrey et al., 1998) and the somatosensory system (Deschênes et al., 2003; Xiao and Xu, 2020). In contrast, if SC spiking is driven by the summation of numerous weak inputs, SC neurons could generate new representations by combining the activity of multiple and diverse RGC types, similar to what has been reported in thalamo-cortical visual circuits (Kremkow and Alonso, 2018; Kremkow et al., 2016; Lien and Scanziani, 2018; Niell and Scanziani, 2021; Reid and Alonso, 1995). Anatomically, the spatial spread of RGC axonal arbors in SC (Hong et al., 2011) would support both of these wiring schemes and therefore it is still unresolved whether SC neurons integrate inputs from only a specific RGC type or whether they sample more broadly from a diverse set of RGC functional pathways. These circuit properties will determine how SC neurons integrate retinal activity *in vivo* and thus revealing the principles underlying the functional organization of the retinocollicular circuitry is central for advancing our mechanistic understanding of how SC neurons process visual stimuli and their role in mediating visually guided behaviors.

A hallmark of the retina is its organization into mosaics (Cook and Chalupa, 2000; Roy et al., 2021; Wässle et al., 1981a, 1981b, 1981c), that are present even in species with poor visual acuity like the mouse (Huberman et al., 2008). In these mosaics, RGCs of the same functional type tile the retina in a quasiregular lattice (Cook and Chalupa, 2000; Devries and Baylor, 1997; Roy et al., 2021; Wässle et al., 1981a) which is thought to reflect optimal and efficient encoding of visual scenes (Field and Chichilnisky, 2007; Roy et al., 2021; Wässle et al., 1981a). If the role of neurons in the SC layers is to provide a precise representation of retinal signals to downstream processing stages (Evans et al., 2018; Lee et al., 2020; Shang et al., 2018), an important prerequisite would be that the structure of retinal mosaics is maintained with single cell precision at the output level of the retina, i.e. the RGC axons in SC, such that spatial resolution is not impaired. While it is established that retinal axons maintain an overall retinotopic organization within the SC (Cang and Feldheim, 2013), by forming dense arbors within specific locations and layers of the superficial SC (Cheng et al., 2010; Hong et al., 2011; Huberman et al., 2009; Martersteck et al., 2017), experimental evidence is lacking for how precise RGC axonal arbors reflect the spatial organization of neighboring RGC somata and dendrites within the retina, thus whether retinal axons are organized in mosaics.

The primary obstacle to understanding the fine scale spatial organization of RGC axonal arbors and the resulting functional connectivity is due to technical difficulties in recording RGC activity and the location of their axons simultaneously with their postsynaptic targets *in vivo*. Synaptic connectivity between progressive stages of sensory processing is typically assessed using topographically aligned recordings of somatic activity in the two regions of interest (Bereshpolova et al., 2020; Lien and Scanziani, 2018; Reid and Alonso, 1995; Usrey et al., 1998). However, this methodology has a low yield of synaptically connected neurons, often restricted to a few pairs recorded simultaneously (Liew et al., 2021; Usrey et al., 1998), which ultimately limits our understanding of how populations of afferent inputs are integrated within target circuits. Here we show that high-density electrodes overcome this technical limitation and that measuring the activity and location of RGC axons simultaneously with their postsynaptic targets in the midbrain at a large scale *in vivo* is possible. Employing this method, we investigate whether RGC axons are organized in mosaics in the midbrain, and elucidate how midbrain neurons functionally integrate those afferent inputs *in vivo*. In addition, we demonstrate that the observed wiring schemes and functional patterns are shared principles across mammals (mouse, *Mus musculus*) and birds (zebra finch, *Taeniopygia guttata*).

## RESULTS

### Recording afferent axons and local neurons simultaneously using high-density electrodes

To study the functional organization of the retinocollicular circuity we used high-density electrodes (Neuropixels probes (Jun et al., 2017)) to record extracellular neuronal activity in the visual layers of mouse SC *in vivo*. We targeted SC with a tangential recording configuration that places hundreds of recording sites within SC (Figures 1C and S1, see Methods). We discovered that the high spatiotemporal sampling of those electrodes, together with their low noise level, allows distinguishing waveforms from somatic action potentials (APs) of SC neurons (Figure 1D, right) from axonal APs of RGC axons (Figure 1D, left; see Figure S2 for waveform classification). Both types of waveforms can be sorted (see Methods) into well isolated single unit clusters with clear refractory periods (Figure 1D, see spike train auto-correlogram “ACG”) and comparable good quality metrics such as AP amplitude and isolation distance (see Figure S3 for details). The majority of waveforms of somatic APs are biphasic and with a small spatial spread (Figure 1D, right). In contrast, the waveforms of afferent axons have a larger spatial spread (Figures 1D, left and S3) and are composed of fast bi/triphasic components caused by the axonal AP and the axons terminal responses (Swadlow and Gusev, 2000) followed by a second slower trough corresponding to the synaptically induced dendritic activity in postsynaptic SC neurons (Figure 1D, left, arrows; Figure S2). Furthermore, we observed that APs propagate along an axonal path in the multi-channel waveform view (Figure 1D, left), with conduction velocities in the range reported from retinal afferents to the SC (Rhoades and Chalupa, 1979) (Figures S1G-I, mean conduction velocity = 3.5±1.3 m/s, n = 283 RGC axons). RGC axons innervate the SC along the anterior-posterior axis (Figures 1A, 1C). Since we could observe the AP propagation only in recordings aligned with the anterior-posterior axis (Figures 1D and S1C-I) but not in recordings aligned with the medio-lateral axis (Figures S1J-M), our observations suggests that the axonal waveforms in our recordings are retinal afferents making synaptic connections onto SC neurons. To confirm this prediction, we performed a series of *in vivo* pharmacological experiments (Figures 1E-1F and S4). We injected Muscimol, a GABAA receptor agonist, into the SC *in vivo* to verify that the triphasic waveforms are signals from long range axons that innervate SC. As expected, these signals were not suppressed by Muscimol application (Figure 1E, green). We then injected a synaptic blocker (see Methods) into the SC to confirm that the second negative waveform component originates from postsynaptic responses in SC neurons (Figure 1F). Finally, we applied tetrodotoxin (TTX) to the eye of the mouse to show that axonal waveforms originate from the retina and do not arise from other sources, e.g. cortex (Figure 1E, light green).

These results demonstrate that the triphasic waveforms recorded in the SC originate from RGC axons making synaptic contacts with SC neurons. In addition, the small distance between recording sites allowed us to capture and isolate activity from neighboring RGCs that are members of the same retinal mosaic (Figures 1G and S5). These RGCs are identified by similar functional responses to a visual chirp stimulus (Baden et al., 2016), non-overlapping receptive field centers, and putative electrical coupling, evident in the double peaks in the cross-correlograms (CCG) which is a defining characteristic of coupling between same type RCGs in the retina (Mastronarde, 1983) (Figures 1G and S5B/C). Being able to record from RGC axons of neighboring RGCs shows that the spatial and temporal resolution of our method is sufficient to reliably isolate individual RGC axons in the SC *in vivo*. Individual recordings yielded a high number of simultaneously recorded RGC axons (∼ 40% of clusters) and SC neurons (∼ 60% of clusters) (n = 27 recordings, n = 1199 RGC axons, n = 1840 SC neurons). Recorded RGC axons and SC neurons covered a large region within the SC circuit and across the visual field (Figure 1H, see Figure S1), with RGC axons deriving from a diversity of functional retinal pathways (Baden et al., 2016) (Figure 1I).

Taken together, we show that high-density electrodes enable recording of the activity and location of afferent axonal APs simultaneously with the AP of post-synaptic targets at a large scale *in vivo*, both in anesthetized and awake mice (Figure S1N). This method thereby permits to study the spatial organization of afferent axons and the resulting functional connectivity with their target neurons *in vivo*.

### Retinal axon mosaics

A precise representation of retinal signals at the level of SC neurons would require that retinal mosaics are mapped isometrically onto the SC surface, i.e. the spatial relationship between RGCs within the retina is reproduced at the level of their axons in SC. However, whether the mosaic structure is indeed maintained with such a precision at the level of RGC axons is an open question. Addressing this issue requires relating the locations of neighboring RGCs from the same mosaic in the retina to the anatomical locations of their axonal arbors within SC *in vivo* (Figure 2A, top). The location of the RGC dendritic arbor can be estimated *in vivo* from the visual receptive field (Figure 2A, see Methods) and the anatomical location of the RGC axonal arbor can be inferred from the RGC waveform on the high-density electrode (Figures 1D/1H). In particular, the recording sites that contain the postsynaptic component of the triphasic RGC waveform (Figures 1D/F and 2A bottom-right) identify the anatomical locations where the RGC axonal arbor makes synaptic contacts onto dendrites of SC neurons. We define this area on the probe as the RGC axonal synaptic contact field (AF) and use this *in vivo* measurement as a proxy for the anatomical location of the RGC axonal arbor within SC. Since the recording sites on the Neuropixels probe are organized in a checkerboard pattern with 480 rows (electrode pitch = 20 µm) and four columns (electrode pitch = 16 µm) it is possible to estimate the position and spatial extent of the AFs both along and across the probe (Figure 2A, see Method).

**Figure 2:**
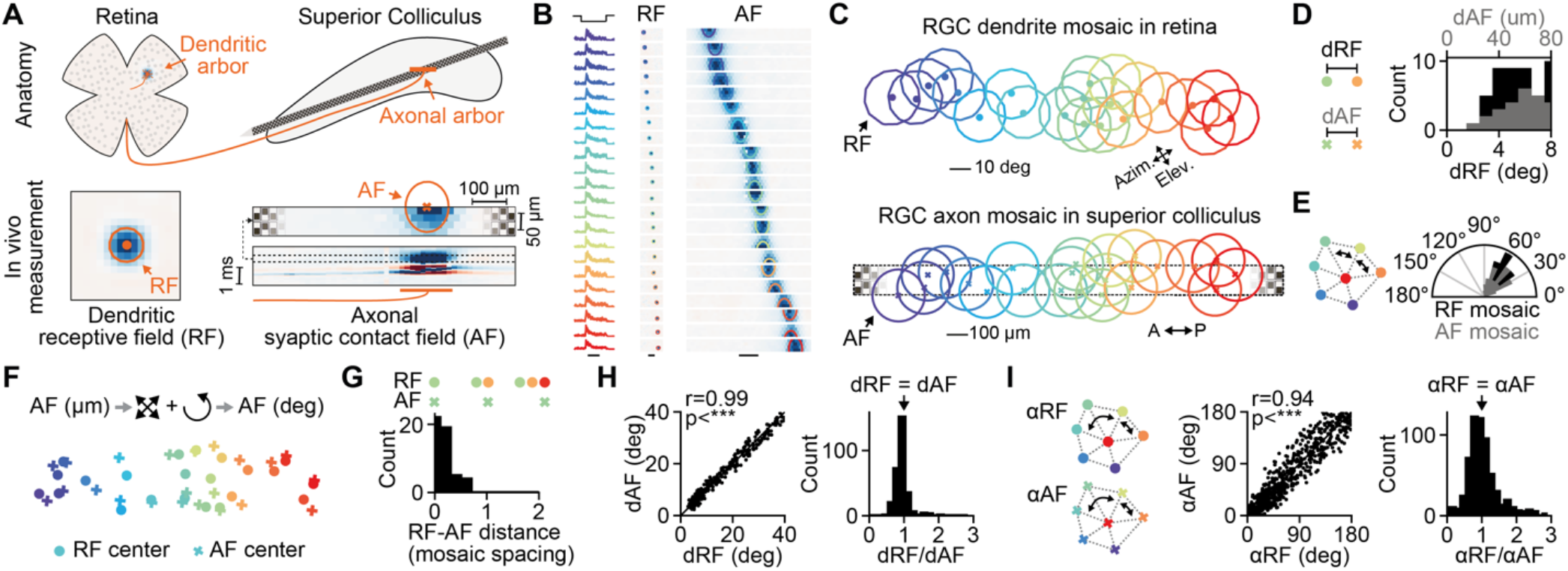
Retinal axon mosaics. (A) Characterization of RGC dendritic and axonal arbors using *in vivo* measurements. The spatial location of the RGC dendrites can be estimated *in vivo* by mapping the visual receptive field (RF), left. The spatial location the RGC axonal arbors in SC can be measured *in vivo* by the axonal synaptic contact field (AF), right. The AF is the area on the high-density electrode with evoked postsynaptic responses in SC. (B) Simultaneous measurement of RGCs belonging to the same functional mosaic. RGC functional type was identified using a chirp stimulus (left, shown are the normalized PSTHs) and receptive field polarity (middle). In this example RGCs were from the OFF_transient_ type. The corresponding RFs and AFs (right) cover a large extend of the visual field and SC tissue. The contours on the AF colormaps show the Gaussian fits to the AF. Scale bars: 1 s, 10 deg, 100 µm. (C) The mosaic of the RGC dendrites in the retina (RF) and the corresponding mosaic of the RGC axons (AF) in SC from data shown in B. Note the similarity between the mosaics at the level of the retina (RF) and axons in SC (AF). (D) Histogram of the distances between RF centers (dRF, bottom axis) and AF centers (dAF, top axis). The gap at close distances is a hallmark of the quasiregular organization of retinal mosaics. (E) Nearest neighbor angles within the RF and AF mosaics revealed by Delaunay and Voronoi tessellations analysis. The peak around 60 deg is indicative of a hexagonal organization. (F) Overlay of the RF/AF centers of the mosaics shown in C. For the overlay the AF mosaic was transformed from SC space (µm) into visual space (deg) by scaling and rotating the AF centers. This transformation preserves the geometrical properties of the AF mosaic. (G) Histogram of the distances between RF and AF centers in the unit of mosaic spacing (n = 50 RGCs, n = 7 mosaics). The mosaic spacing is the median RF distance between neighboring RGCs. (H) Distances between RGCs in the RF and AF mosaics are similar. dRF plotted against dAF and the histogram of the ratio between dRF and dAF (n = 294 RGC pairs). (I) Angles between the RGCs in the RF and AF mosaics match. Angles between triples of RGCs measured in the RF and AF mosaics (n = 627 RGC triples) and the histogram of the ration αRF/αAF. Note, in E we show the angles between nearest-neighbors to test for hexagonality. In I we show the angles across larger distances to estimate the overall geometrical similarity between the RF and AF mosaic. For all panels: *** = p < 0.001.

To investigate how the axonal arbors of neighboring RGCs of the same type organize in SC requires identifying RGCs belonging to the same functional mosaic. Three criteria can be used to identify RGC from the same mosaic: 1) comparing the visually evoked activity of RGCs to a chirp stimulus (Figures 1G and 2B, left); 2) comparing the polarity (ON or OFF) of the RGC visual receptive fields (Figure 2B, middle; see Methods), and 3) identifying putative electrically coupled RGCs via spike train cross-correlation (Figures 1G and S5B). High similarity in the evoked chirp responses as well as same polarity of the receptive fields indicate that RGCs are of the same functional type while putative electrical coupling further increases the confidence that these RGCs belong to the same retinal mosaic (Mastronarde, 1989) (Figure S5). However, it is important to note that not observing electric coupling does not indicate that RGCs are from different mosaics. Having sorted recorded RGCs into functional types we studied and compared the spatial organization of their mosaics at the level of the retina using the RFs and at the level of the axon in SC using the AFs.

Figure 2B shows a recording in which we were fortunate to capture a large number of RGCs from a similar functional type, OFF-transient in this case (Figure 2B, left, n = 22 RGCs). The AF positions of the RGCs gradually changed along the probe and within SC (Figure 2B, right) with the corresponding RF locations varying in elevation and azimuth (Figure 2B, middle). Overlaying the RF of all RGCs confirmed well-known properties of retinal RF mosaics: RF centers were non-overlapping, quasiregular, and hexagonally arranged (Figure 2C, top; color-code same as in Figure 2B). Measuring the distance between RF centers (dRF) showed a clear gap at close distance (Figure 2D, black), indicative of non-overlapping RF centers. Estimating the angles between nearest neighbors in the RF mosaic, employing the Delaunay and Voronoi tessellations (Zhan and Troy, 2000) (see Methods), showed a peak around 60 deg (Figure 2E, black), which is characteristic for hexagonal organizations. Both measurements thus confirm that our method can capture RGCs belonging to the same RGC mosaic *in vivo* (Figure S5).

Remarkably, the mosaic organization in the retina was almost perfectly preserved at the level of the RGC axons within SC (Figure 2C, compare top and bottom). As for the RFs, the histogram of the distances between AF centers showed a clear gap at close distance (Figures 2D and S7C, gray) and the AF centers were quasiregular and hexagonally arranged (Figure 2E, gray), revealing that RGC axonal arbors are organized in mosaics (Figures 2C and S5G). To assess the geometrical similarity between the RF and AF mosaics we transformed the AF mosaic from the anatomical space in SC (µm) into the visual space of the RFs (deg), by linearly scaling and rotating the AF mosaic to match the size and orientation of the RF mosaics (Figure 2F, see Methods). This transformation preserves the spatial relationship between AF centers and allows to quantify the similarity between the RF and AF mosaics. To quantify the similarity, we measured the distances between RF and AF centers of individual RGCs and divided these values by the mosaic spacing in each mosaic (mosaic spacing = median nearest neighbor’s RF distance). Our results show that RF and AF centers closely overlay and align and that the RF and AF mosaics are geometrically similar (Figures 2G, median distance 0.26±0.17 mosaic spacing, n = 50 RGCs from n = 7 mosaics and n = 5 mice). To further characterize the similarity, we compared the RF and AF distances between RGC pairs (Figures 2H, median dRF/dAF = 1.00±0.18, n = 294 RGC pairs from n = 7 mosaics and n= 5 mice) and compared the angles within the RF and AF mosaics (Figure 2I, median αRF/αAF = 1.04±4.76, n = 627 angles from n = 7 mosaics and n = 5 mice). All these measurements show a close correspondence between the RF and AF centers and strongly suggest that RGC axon mosaics within SC are almost perfect copies of the RGC dendritic mosaics in the retina. Thus, on the level of single cells, RGC axons provide precise isomorphic representation of the retina as input to SC. But how do SC neurons sample from this precisely organized afferent input? To answer this question, we next studied monosynaptically connected RGC-SC pairs.

### Measuring monosynaptic connectivity *in vivo* at a large scale

Simultaneous recordings of RGC axons and SC neurons permit identification of synaptically connected RGC-SC pairs. To assess synaptic connectivity, we employed established cross-correlation analysis methods (Bereshpolova et al., 2020; Reid and Alonso, 1995) (Figures 3A/B and S6, see Methods). Connected RGC-SC pairs were identified in the cross-correlograms by significant transient and short-latency increases in the spiking probability (Figures 3A and S6; peak latency = 1.54±0.38 ms, n = 1048 connected pairs), a hallmark of monosynaptic connectivity in vertebrate nervous systems (English et al., 2017; Jouhanneau et al., 2018; Lien and Scanziani, 2018; Reid and Alonso, 1995; Usrey et al., 1998). Unconnected pairs do not show transient peaks (Figures 3B and S6B/C). Depending on the number of recorded RGC axons, we could identify up to 229 monosynaptic connections in individual recordings (Figures 3C and S6), yielding in total above one thousand measured connected RGC-SC pairs across multiple experiments (Figure 3D, n = 1048 connections, n = 1199 RGC axons, n = 32 experiments, average probability of detecting RGC-SC connections per RGC axon = 87%). This high number of measured connected pairs results from the close proximity of RGC axons and SC neurons on the high-density electrode probe (Figure 3E, left; distance within SC = 70.63±141 µm, n = 1048 connected pairs) and similar receptive field locations (Figure 3E, right, RF distance = 4.64±4.75 deg, n = 530 connected pairs). Due to the large number of identified connections, we were able to identify diverging connections from single RGC onto multiple SC neurons (Figure 3F, left) and converging connections from multiple RGCs onto single SC neurons (Figure 3F, right), permitting us to investigate the functional organization of the retinocollicular circuit *in vivo*.

**Figure 3:**
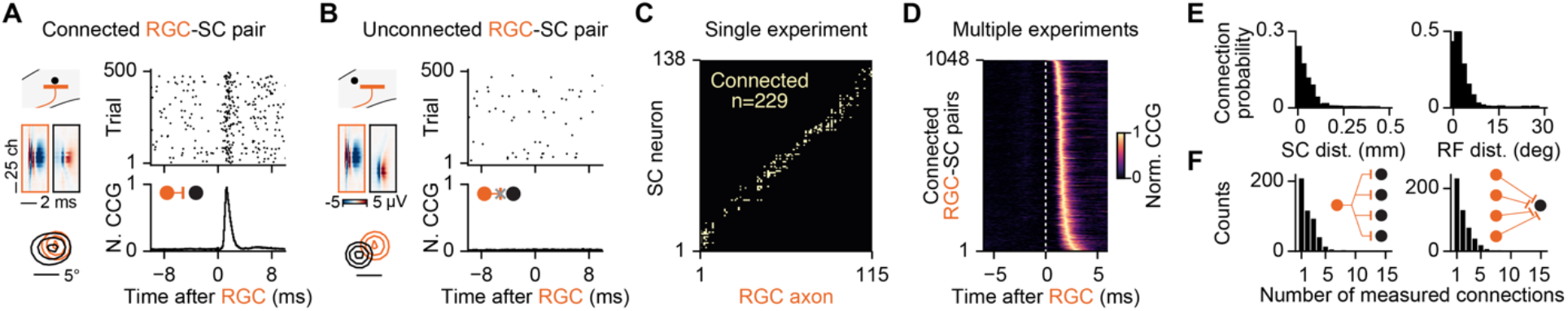
Measuring afferent monosynaptic connections *in vivo* at a large scale. (A) Examples of a monosynaptically connected RGC-SC pair. Top, raster plot of SC spiking activity triggered on RGC spike times. Bottom, cross-correlogram (CCG) between the RGC and SC spiking activity. The short latency peak in the CCG is a hallmark of synaptic connections in vertebrates. Note the close distance of the RGC axon and SC neuron waveforms on the probe (middle) and the overlapping RFs (bottom). (B) Unconnected RGC-SC pair. Unconnected pairs do not overlap in SC and visual space. (C) Connectivity matrix between RGC axons and SC neurons recorded in a single experiment. Yellow marks identify connected pairs. RGC axons and SC neurons are sorted by their location in SC. (D) CCGs of connected pairs across multiple experiments, sorted by peak latency. (E) Connection probability as a function of SC distance and RF distance. (F) Number of measured RGC-SC connections per RGC axon (left) and SC neuron (right).

### Synaptic organization of the retinocollicular circuit *in vivo*

Previous studies have shown that single RGC spikes reliably trigger postsynaptic activity in neurons of the visual thalamus, dorsal lateral geniculate nucleus (dLGN) (Kaplan and Shapley, 1984; Usrey et al., 1998), and that the majority of dLGN spikes are driven by RGC activity (Usrey et al., 1999). It is unknown, however, whether this strong drive and coupling are common principles of RGC connections and are therefore also present in the retinocollicular pathway (Figure 1B, top), or whether the retinocollicular circuit is different such that SC neurons receive weak inputs from numerous RGCs (Figure 1B, bottom). To differentiate between these distinct modes of signal transmission, we examined the activity of connected RGC-SC pairs (Figure 4). Our data shows that individual RGC APs can trigger responses in the postsynaptic SC neuron (Figure 4A, “1”) or fail to be transmitted (Figure 4A, “2”), and that SC APs can occur without input from that specific RGC (Figure 4A, “3”). To quantify these observations, we estimated the connection efficacy and connection contribution (Usrey et al., 1999) for each connected pair. The efficacy is the probability that an RGC input triggers an AP in the postsynaptic SC neuron (Figure 4B, left). In the example shown in Figures 4A-B, the efficacy was ∼ 17% and across the population, we observed a log-normal distribution of connection efficacies, with a few very strong connections up to ∼ 50% efficacy, but primarily weaker connections (Figure 4C, efficacy = 4.01±4.20%, maximum efficacy = 48.08%, n = 1048 connected pairs). Next, we estimated the connection contribution, which characterizes the fraction of SC APs that are driven by RGC activity from a single RGC unit and therefore provides a measure for how strong SC neurons are coupled to the activity of individual RGC inputs. High contribution values indicate that SC neurons are primarily driven by their RGC afferent inputs while low contribution values reflect that SC neurons are driven by inputs from other sources. Our data show that SC neurons can be strongly coupled to retinal inputs, such that a large fraction of SC APs are preceded by retinal APs (Figure 4B, right), however we primarily observed weaker coupled pairs (Figure 4D, contribution = 15.31±11.36, maximum contribution = 78.63, n = 1048 connected pairs). Across the population, we discovered a log-normal distribution of connection and coupling strengths. Log-normal distributions of connection strength are widely observed in the vertebrate brain (Buzsáki and Mizuseki, 2014; Cossell et al., 2015; Jouhanneau et al., 2015, 2018), including human (Campagnola et al., 2021), which could be the result of circuit refinement during development (Dhande et al., 2011) by which only a few strong connections remain after the refinement process. Our data support this view whereby RGCs establish only one or a few strong connections with SC neurons and multiple weaker connections (Figures 4E/F and S6E). Such sparse and strong connections are optimal for reliably transmitting signals along signaling pathways (Kumar et al., 2010) and suggest that the retinocollicular circuitry is optimally wired for transmitting retinal activity in a functional specific manner.

**Figure 4:**
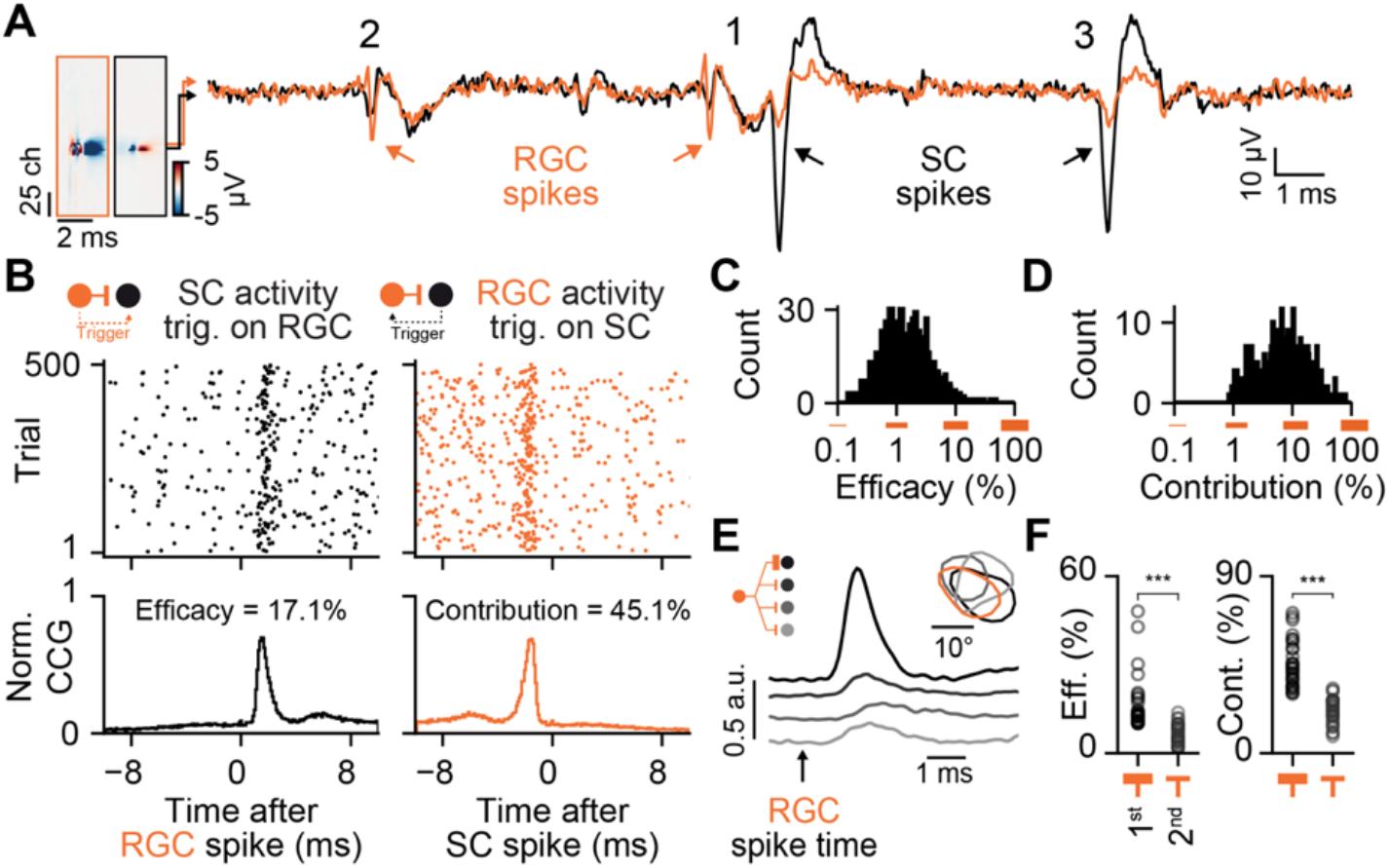
SC neurons are strongly driven by retinal afferent inputs. (A) Example traces showing the electrical signals of a monosynaptically connected RGC axon (orange) and SC neuron (black) pair. 1) the RGC spike triggers a SC spike, 2) failed transmission, 3) SC spike without RGC input. (B) Strong RGC-SC coupling. Left, SC spiking activity relative to the RGC spike times of the example pair shown in a. Note that the RGC input strongly drives spiking in the connected SC neuron, estimated by the connection efficacy. Right, a large fraction of SC spikes are driven by RGC activity, estimated by the connection contribution. (C) Population histogram of efficacy values (n = 1048 connected pairs). Note the log-normal distribution, with few strong and many weak connections. (D) Contribution of all measured connections (n = 1048 connected pairs). (E) Example of a divergent connection with one strong and several weak connections. Inset shows the receptive field contours of the recorded neurons. (F) Efficacy and contribution measurements for the strongest and second strongest connection (efficacy: 1^st^ = 16.52±9.08%, 2^nd^ = 6.56±3.29%, p < 0.001; contribution: 1^st^ = 45.22±11.77%, 2^nd^ = 21.53±6.36%, p < 0.001; n = 30 divergent connections).

### Functional organization of retinal afferent connections in SC

Next, we studied the functional organization of the retinal inputs to SC neurons as it is still largely unknown how SC neurons integrate convergent inputs from the diverse set of retinal functional pathways (Ellis et al., 2016). To this end, we characterized and compared the visually driven responses of connected RGC-SC pairs. We started to investigate the retinotopic alignment and functional similarity of connected RGC-SC neurons by measuring the spatiotemporal receptive field (STRF) using a sparse noise stimulus (Kremkow et al., 2016) that reliably triggers spatiotemporally organized visually evoked activity, in both RGC and SC neurons (Figure 5A, see Methods). Comparing the STRFs of connected neurons provided a first measure of their functional similarity. The similarity (S) was estimated by the correlation coefficient between the STRFs of connected pairs. A value of 1 corresponds to STRFs that are perfectly correlated, a value of 0 reflects uncorrelated STRFs. Our data reveal that RGC-SC pairs with more similar STRFs are more strongly connected (Figure 5A, compare black-gray pair vs. orange-black pair), a relationship also found across the population of connected RGC-SC pairs (Figure 5B). Another important functional property of visual neurons is their selectivity to the direction of motion. Previous work indicates that direction-selective SC neurons receive directionally tuned RGC inputs (Shi et al., 2017). However, a confirmation on the level of monosynaptically connected direction-selective RGC-SC pairs has remained pending. We compared the preferred directions (PD) of monosynaptically connected RGC-SC pairs which confirm that connected and direction-selective RGCs and their SC targets have similar preferred directions (Figures 5C/D, see Methods).

**Figure 5:**
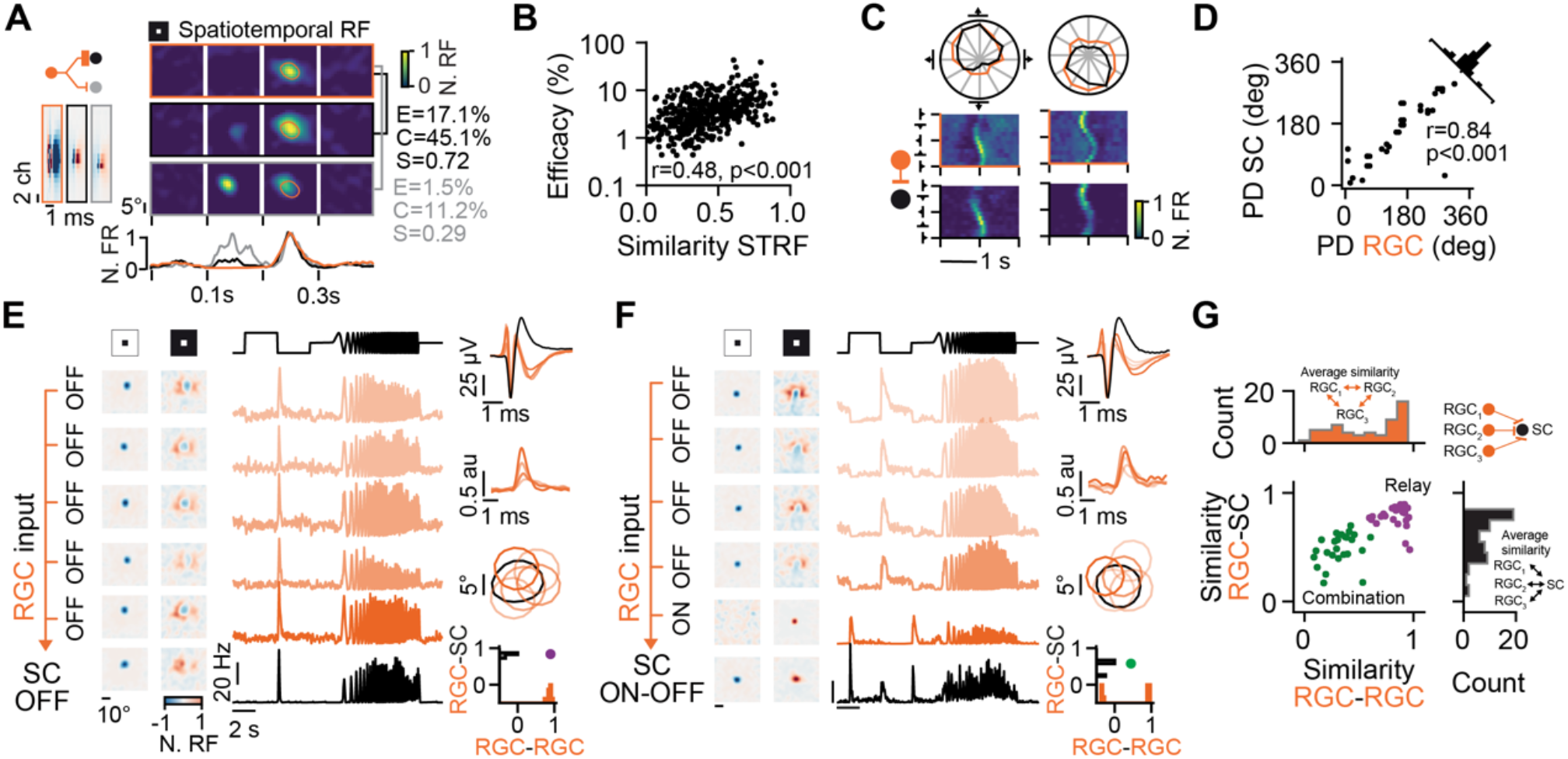
Functional organization of the retinal afferent connections. (A) Example of a divergent connection between an RGC (orange) and two of its postsynaptic targets in SC (black and gray). Shown are the spatiotemporal receptive fields (STRF, top), and the temporal response at fine temporal scale (bottom). E = efficacy, C = contribution, S = similarity of the STRFs. (B) Correlation between the similarity of STRFs and connection efficacy (n = 530 connected pairs). (C) Directional tuning of connected RGC-SC pairs. Top, polar plot indicating overlapping directional tuning of connected pair (orange and black). Bottom, normalized firing rate in response to moving bars. (D) The preferred directions (PD) of RGC axons and their connected postsynaptic SC target neurons match (mean PD difference = 24.23±29.15°, n = 50 connected pairs). (E) Relay motif example of an SC neuron receiving convergent inputs from a pool of RGCs with similar functional responses. Left, receptive fields. Middle, responses to the chirp stimulus. Right, spike waveforms, CCGs, contours of RFs, histogram of similarities between RGC-RGC (orange) and RGC-SC (black). (F) Combination motif example, same format as e. Both ON and OFF RGCs converge onto the SC neuron. (G) Functional convergence, population analysis. Top, histogram of the average RGC-RGC correlations of convergent pools. Note that some afferent input pools are highly correlated with values close to 1 while others convey a mixed input with values close to 0.5. Right, histogram of the average RGC-SC correlation. Scatter plot of the average RGC-RGC similarity vs. the average RGC-SC similarity for SC neurons with >= 3 RGC inputs. Note both relay motifs (violet) and combination motifs (green) across the population of SC neurons (n = 57 SC neurons, n = 27 recordings).

These results support the notion that the retinocollicular circuit is organized in a functionally specific manner, with some response features of RGC stimulus selectivity being passed on onto their respective strongest connected SC neurons. However, not all RGC types are direction selective and the distribution of similarity values from dissimilar (S = 0) to identical (S=1) (Figure 5B), suggests that not all SC neurons receive convergent inputs from a functionally homogenous pool of RGCs, raising the question whether different functional convergence motifs exist. To clarify this issue, we extended our functional characterization by analyzing responses to a chirp stimulus that allows classifying RGCs into different functional types (Baden et al., 2016) (Figure 1I, see Methods). Studying the convergence of RGC axons to single SC neurons revealed that relay and combination motifs exist (Figures 5E/F). In relay motifs, a SC neuron receives convergent inputs from a population of RGCs of the same functional type, such that the evoked responses were similar to the presynaptic RGC pool (Figure 5E, note the similarity between receptive fields as well as evoked chirp responses of RGCs and the SC neuron). In contrast, in combination motifs SC neurons pool afferent inputs from a few but diverse set of functional RGC types (Figure 5F). We characterized the functional specificity of the convergence motifs by calculating the similarity (correlation) of the chirp responses among the presynaptic RGCs (Figures 5E/F, bottom-right, orange) and the similarity between the RGCs and the postsynaptic SC neuron (Figures 5E/F, bottom-right, black; see Methods). Values close to 1 for both measurements reflect a relay motif (Figure 5E) while lower values indicate a more diverse functional relationship (Figure 5F) between the presynaptic RGCs and the postsynaptic SC neuron. Across the population of SC neurons, the functional RGC-SC convergence follows at least two motifs with presynaptic RGC pools being homogenous (Figure 5G, violet) or functionally diverse (Figure 5G, green).

### Comparing the mammalian and avian retinotectal circuit

Our data demonstrate that the synaptic and functional organization of the retinocollicular pathway of the mouse is characterized by sparse and strong connections that establishes functional specific wiring between retina and SC. Since the retinofugal pathway to the midbrain is highly conserved across all classes of vertebrates (Basso et al., 2021), we hypothesized that these wiring principles are of general nature and therefore likely to be found in non-mammalian vertebrate species, e.g. birds. To test this hypothesis, we studied the synaptic and functional organization of retinal afferent inputs to neurons in the optic tectum of the zebra finch employing our high-density electrode method to simultaneously measure RGC axons and connected OT neurons *in vivo* (Figures S7/8). Our data confirms that RGC axons form mosaics in the OT of the zebra finch that precisely reflect the mosaic at the RF level in the retina (Figures 6B/C and S7G-K, finch: median dRF/dAF = 1.01±0.28, n = 55 RGC pairs; median αRF/αAF = 1.04±1.14, n = 155 angles). Similar to the mouse SC, zebra finch OT neurons receive a limited pool of RGC afferents (Figures 6D and S8F) with a log-normal distribution of RGC connection strengths (Figures 6D and S8B). As a consequence, OT neurons are strongly driven and tightly coupled to their RGC afferent inputs (Figures 6E and S8, finch: efficacy = 7.16±7.16%, maximum efficacy = 39.26%, contribution = 13.36±11.73%, maximum contribution = 58.83%, n = 105 connected pairs), which establishes a precise retinotopic representation of retinal activity in OT/SC neurons (Figure 6F) and functional specific wiring (Figures 6G and S8D, r = 0.69, p > 0.001, n = 53 connected pairs). Despite the higher spatial resolution of the avian visual system (Schmidt et al., 1999) and the large evolutionary distance between mammals and birds, our results indicate that retinal afferents are integrated by zebra finch OT neurons following similar principles to neurons in the mouse SC (Figures 6 and S7/8). Taken together, our data strongly support the notion that retinotectal circuit follows similar wiring principles across vertebrate species.

**Figure 6:**
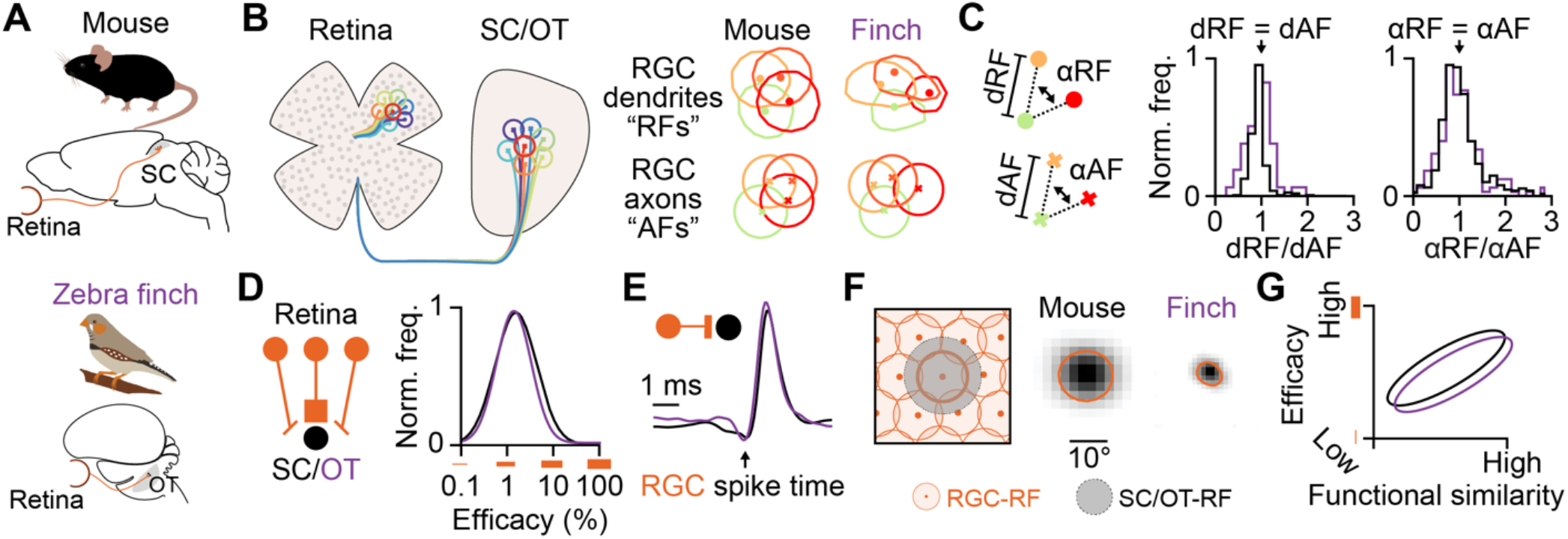
Wiring principles of the retinotectal circuit in mammals and birds. (A) Retinotectal pathway in mammals and bird. The mouse superior colliculus (SC) and the zebra finch optic tectum (OT) receive direct inputs from retinal ganglion cells (RGC), the output cells of the retina. (B) Retinal axons are organized in mosaics in SC/OT that reflect the mosaic organization in the retina. Shown are examples of a RGC RF and AF mosaic from mouse and finch. (C) The RGC axon mosaics provide a single cell precise isomorphic representation of the RGC RF mosaics as input to the SC/OT, evident by the similarity in distances and angles between RF and AF centers. Distances: n = 294 RGC pairs in the mouse and n = 92 RGC pairs in the finch. Angles: n = 627 RGC triples in the mouse and n = 199 RGC triples in the finch). Note, the mouse data is also shown in Figures 2H/I and shown here again to allow a comparison with the finch data. (D) Limited convergence in the retinotectal circuit. RGC provide sparse but strong inputs to SC/OT neurons, established by a log-normal distribution of connection efficacies in both vertebrate species. (E) Individual RGC inputs strongly drive SC/OT spiking. Average cross-correlograms of connected RGC-SC pairs (n = 1048 connected pairs) and of connected RGC-OT pairs (n = 105 connected pairs). (F) Precise retinotopic convergence. Average receptive field of SC/OT neurons (grayscale colormap) and their RGC afferents (orange contour) (n = 444 RGC-SC pairs; n = 47 RGC-OT pairs). (G) Schematic of the functional specificity of RGC-SC and RGC-OT connections. See Figures 5B and S8D for experimental results.

## DISCUSSION

### Large scale paired recordings with high-density electrodes *in vivo*

We have identified a novel approach to study the functional connectivity across brain regions *in vivo* by using high-density electrodes recordings to capture the electrical activity of both afferent axons and the somatic activity of local neurons simultaneously on the same probe. Revealing the functional composition of presynaptic pools (Wertz et al., 2015), including afferent connections, has been technically challenging, as it requires recording with multiple carefully aligned electrodes in the afferent and target brain region (Bereshpolova et al., 2020; Lien and Scanziani, 2018; Liew et al., 2021). High-density electrodes solve this issue by recording the activity of afferent axons and their postsynaptic target neurons simultaneously on nearby channels on the same probe (Figures 1D and 3A/E), thereby yielding an unprecedentedly large number of connected pairs *in vivo* (Figures 3C/D). The continuously increasing number of recording sites and reduction in site-to-site distance (Steinmetz et al., 2021) of high-density electrodes, together with a growing applicability of these probes for chronic recordings, will open up the possibility to study the functional connectivity between connected brains regions in freely behaving animals.

### Synaptic contact fields of afferent axons

Revealing the functional properties of afferent axons and how those axons organize within target structures is central for reaching a mechanistic understanding of functional neural circuits. Our novel method of measuring the axonal synaptic contact field of afferent axons, on the two-dimensional layout of the high-density electrode *in vivo*, opens up opportunities to investigate the principles of how afferent inputs organize in other parts of the brain, e.g. how thalamic afferents organize within sensory cortices (Jin et al., 2011; Kremkow and Alonso, 2018; Kremkow et al., 2016). Furthermore, our work demonstrates that it is feasible to measure and characterize sub-compartments of afferent connections *in vivo*, including synaptically evoked dendritic responses in postsynaptic neurons (Swadlow and Gusev, 2000), on a single high-density electrode (Figure 1D). Previous work using spike triggered field potentials revealed important insights about the functional and synaptic properties of thalamocortical (Bereshpolova et al., 2020; Jin et al., 2011; Stoelzel et al., 2008; Swadlow and Gusev, 2000; Swadlow et al., 2002) and corticocollicular circuits (Bereshpolova et al., 2006). However, these studies required technically challenging paired recordings with well aligned electrodes in both brain regions. Our approach overcomes this technical challenge by measuring the action potential and location of the afferent axon together with the evoked synaptic field simultaneously on the same probe. Moreover, recent work using planar high-density electrodes showed that it is possible to characterize connectivity and function using extracellular electrodes in *ex vivo* preparations (Shein-Idelson et al., 2017). Our study now shows that high-density electrodes can be used for studying synaptic physiology of afferents *in vivo*, which opens up new avenues in how to probe both function and structure of neural circuits; including the plasticity of afferent synaptic inputs. Finally, high-density electrodes could potentially be used to refine models of the origin of local field potentials (Hagen et al., 2017) by relating afferent synaptic inputs to ongoing local field potentials in cortical and sub-cortical brain structures in future studies.

### Retinal axon mosaics

Retinal mosaics are one of the most spatially precise organizations in the vertebrate brain (Field and Chichilnisky, 2007; Roy et al., 2021; Wässle et al., 1981a). However, whether the mosaic structure is unique to the input level of the retina (RGC dendrites) or also maintained at the output level or the retina (RGC axons) has been an open question since the discovery of retinal mosaics more than 170 years ago (Cook and Chalupa, 2000; Hannover, 1843). Our experimental data show an extraordinary correspondence between the spatial organization of the retina at its input (dendrites) and its output (axons). Retinal axons form mosaics that isomorphically and with single cell precision represent the spatial organization of the retina as input to the superficial layers of the midbrain. While the isomorphic representation of the retinotopic map on a larger scale is a known hallmark of the visual system (Cang and Feldheim, 2013), the single cell precision of this mapping at the level of the RGC axons in the midbrain has not been shown before. Furthermore, while RGC axons innervate the visual layers of the mammalian SC from below (Cang and Feldheim, 2013; Huberman et al., 2008), RGC axons in the avian visual system grow into the OT from the outside of the most superficial layers (Schmidt et al., 1999). Despite these anatomical differences on the macroscopic level, the fine scale organization of the RGC axons within the target layers of the midbrain and the resulting functional connectivity appears to be comparable between mammals and birds. This strongly suggests that the highly precise wiring of the retinotectal circuity that we discovered is essential for visually guided behaviors in vertebrates. It also raises the question what developmental mechanisms underlie this single cell precise mapping between the retina and the midbrain and whether this precision is unique to vision or a general principle of how sensory afferents organize in the midbrain (Benavidez et al., 2021; Drager and Hubel, 1976). Finally, sensory cortices are topographically organized (Kremkow and Alonso, 2018) and identifying similarities and differences in the mapping of afferents into the midbrain and cortex will provide important insights into the role these circuits play for mediating perception and behavior.

### Functional organization of the retinotectal circuit

We characterized the functional organization of the retinotectal connections in both the mammalian and avian visual systems. Our results show that retinotectal wiring is governed by limited functional convergence, precise retinotopic and functional specificity and strong efficacy (Figure 6). The strong drive from RGC axons onto SC/OT neurons, together with the limited convergence, suggests that a major role of the retinotectal connections is to provide a faithful representation of the visual world to postsynaptic first order SC neurons, so that downstream neurons and circuits (Evans et al., 2018; Lee et al., 2020; Reinhard et al., 2019; Shang et al., 2018), have a reliable access to the sensory environment encoded in diverse retinal pathways (Baden et al., 2016; Savier et al., 2019). This efficient way of integrating afferent inputs is reminiscent to the way neurons in the visual thalamus (dLGN) integrate retinal inputs (Rosón et al., 2019; Usrey et al., 1999), but it is contrasting to the thalamocortical system in which thalamic afferent inputs to excitatory neurons in cortex are weak (Bruno and Sakmann, 2006; Lien and Scanziani, 2018; Reid and Alonso, 1995). Thus, the main brain regions involved in visual processing, midbrain and visual cortex, integrate their afferent inputs in different ways, suggesting distinct roles in sensory processing and visually guided behaviors.

We also observed combination mode motifs and it is still unresolved what determines whether an SC/OT neuron receives functionally specific or diverse afferent inputs. One possibility is the downstream targets of the postsynaptic SC/OT neuron (Reinhard et al., 2019). Another possibility is the location within the SC/OT circuit, or the cell type of the SC/OT neuron since it is known that excitatory and inhibitory neurons in visual cortex receive different types of functional convergence from thalamus (Alonso and Swadlow, 2005; Bereshpolova et al., 2020). Despite the evidence that SC neurons can inherit directional tuning from their RGC afferents (Shi et al., 2017), several response properties of SC neurons cannot be explained by simple inheritance from the retina. For example, SC neurons can be orientation selective (Ahmadlou and Heimel, 2015; Feinberg and Meister, 2015; Ito et al., 2017; Wang et al., 2010) with a wide range of orientation preferences present across the population (Ahmadlou and Heimel, 2015; Wang et al., 2010). While some orientation tuned RGC types have been reported, those RGCs mainly encode the cardinal orientations (Nath and Schwartz, 2016) and thus, inheritance alone cannot fully explain orientation preference in SC. A potential additional mechanism could be the convergence of untuned ON- and OFF-center RGCs at the level of SC neurons, similar to the way cortical neurons generate orientation preference from the convergence of ON- and OFF-center dLGN neurons (Jin et al., 2011; Kremkow et al., 2016). However, this hypothesis needs to be thoroughly investigated in future work.

Neurons in vertebrate visual circuits, including SC, are often organized in functional maps (Ahmadlou and Heimel, 2015; Feinberg and Meister, 2015; Kremkow and Alonso, 2018; Li et al., 2020) (but see (Chen et al., 2021)) that are thought to arise, at least in part, due to the spatial organization of the afferent inputs. While a number of studies have addressed how thalamic afferents shape the functional organization of the primary visual cortex (reviewed in (Kremkow and Alonso, 2018)) it remains an open question how the functional maps in SC relate to the spatial organization of the retinal afferents. Modeling work suggests that the moir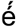 interference between ON- and OFF-center RGCs mosaics (Paik and Ringach, 2011) plays a major role in establishing the functional maps in the thalamocortical visual system. While experimental evidence supporting this hypothesis is still sparse, in large part due to the technical inability to map the organization of retinal mosaics via the thalamus to primary visual cortex *in vivo*, the high-density electrode approach would allow to investigate how RGC mosaics relate to the functional map in SC.

## SUMMARY

In summary, we show that the retinotectal circuit in both mammals and birds is characterized by limited functional convergence with strong and specific connections, established in log-normally distributed connection strength. This precise functional wiring is made possible by single cell precise isomorphic mapping of retinal mosaics to the axonal input level in the midbrain. Because the functional organization of the retinotectal circuit is similar in mammals and birds and resembles the organization principle of retinal inputs to visual thalamus in mammals (Kaplan and Shapley, 1984; Rosón et al., 2019; Usrey et al., 1998), we conclude that retinofugal connections follow a canonical wiring pattern that provides a precise and reliable representation of the visual world to neurons across the different targeted regions in the vertebrate brain.

## ACKNOWLEDGEMENTS

We thank J-M. Alonso, J. Poulet, B. Judkewitz for helpful discussions and materials exchanges during the project; P. Wisinski-Bokiniec, C.J. Whitmire and J-M. Alonso for comments on the manuscript; T. Leva, and P. Schnepel for help with the Neuropixels recordings; Fabian Heim for zebra finch histology; J. Siegle and D. Denman for help with software (OpenEphys) and setup; and J. Colonell for Neuropixels hardware explanations. We thank the whole Neuropixels community for their equipment and support and the Allen Institute for Brain Science for fostering high quality databases. This work was supported by the DFG Emmy-Noether grants KR 4062/4– 1 (JK) and VA 742/2 (DV), the ERC-2017-StG - 757459 MIDNIGHT (DV) and Project number 327654276 – SFB 1315.

## AUTHOR CONTRIBUTIONS

J.S. and J.K. conceived and designed the study; J.S., C.G., J.B., D.V. collected the data; J.S., C.G., H.B., T.L., K-L.T. and J.K. analyzed the data and J.S. and J.K. wrote the manuscript with inputs from all authors.

## DECLARATION OF INTERESTS

The authors declare no competing interests.

## MATERIALS AND METHODS

### Animals, surgery, and preparation

All experiments were pursued in agreement with the local authorities upon defined procedures (LAGeSo Berlin - G 0142/18 and Regierungsprä sidium Oberbayern - ROB-55. 2-2532. VET_02-18-182). During all experiments, maximum care was taken to minimize the number of animals used and their discomfort. *Mice*: Adult male mice (C57BL/6J) from the local breeding facility (Charité-Forschungseinrichtung für Experimentelle Medizin, n = 70) and Charles-River Germany (n = 25) were used. Induction was achieved with isoflurane (2.5% in oxygen Cp-Pharma G227L19A). Once anesthetized, the surgery was performed in a stereotactic frame (Narishige) with a closed-loop temperature controller (FHC-DC) for monitoring the animal’s body temperature. The isoflurane level was gradually lowered during surgery (0.7-1.5%) while ensuring a complete absence of vibrissa twitching or responses to tactile stimulation. During surgery, the eyes were protected with eye ointment (Vidisic). For awake mouse recordings, the head post was implanted two weeks before the recording day and protected with silicone elastomer sealant Kwik-Cast (WPI Germany). Metamizole (200 mg/kg, Zentiva-Novaminsulfon) was administered in drinking water after head post implantation for a recovery period of 3 days. After recovery, the animals were gradually habituated to the recording setup. Craniotomy was performed on the day before the recording. *Zebra finches*: Adult male zebra finches (>180 days post-hatching) were obtained from the local breeding facility at the Max Planck Institute for Ornithology in Seewiesen (n = 7). Birds were anesthetized with isoflurane (1-3% in O2) and head-fixed in a stereotactic instrument (Kopf) while the body temperature was maintained at 40 °C with a homeothermic monitoring system (Harvard Apparatus) with the head tilted by 45 deg to the azimuthal plane. For all experiments, a dental cement-based crown (Paladur, Kuzler) was used to fix the head post, and grounding which settings had to be planned for each desired implantation’s angle to maximize room for probe manipulations. *Recordings*: On the recording day for both, awake (mice) and anesthetized (mice and zebra finches) recordings, the probe was lowered (>4 mm) in the target region according to the stereotactic coordinates, followed by a small withdrawal of 20 to 50 µm to release accumulated mechanical pressure. Once the probe was positioned it was allowed to settle for ∼ 10-20 minutes, receptive fields (RF) were mapped based on their multi-unit-activity (MUA) to confirm that the visual stimulus covered the retinotopic positions of the recorded neurons. Therefore, it was crucial to align the stimulus center to the pupil resting position i.e., 64 deg lateral to the nose of the mouse (Sterratt et al., 2013) and 62 deg lateral to the beak of the finch (Bischof, 1988). Once the visually driven activity was obtained on at least 50 channels (Figure S1), the data acquisition was started and visual stimuli were presented. *Histology*: For histological confirmation and reconstruction of the electrode track, the probe was removed and re-inserted in the same location coated with DiI (Abcam-ab145311) diluted in ethanol. The animal was then sacrificed either with isoflurane (>4%) or a subcutaneous injection of a Ketamine-Xylazine mix (Ketamidor 1 g/mL, Rompun 2%). Cardiac perfusion was performed with phosphate buffer saline solution (PBS) followed by 4% paraformaldehyde (PFA) in PBS. The brains were post-fixed overnight in 4% PFA and stored in PBS until histological slicing was performed using a vibratome (Leica VT1200 S) and the slices were mounted in DAPI-Fluoromount-G (70-100 µm slices, Biozol Cat. 0100-20). Perfused zebra finch brains were transferred to 15% sucrose in PBS for 24 hours post-fixation, and once no longer buoyant, they were moved into 30% sucrose until sinking point. Optic tectum was sliced into 90 µm sagittal sections and mounted using DAKO (Agilent).

### Electrophysiological recordings

Neuropixels probes, Phase 3a and Phase 3B1 (Jun et al., 2017), were used with Open Ephys software (www.open-ephys.org) on either the phase 3A system or the PXIe system (National Instrument NI-PXIe-1071). The signal was amplified and stored in both the local field potential band (LFP, high pass-filtered 0-300 Hz) or the action potential band (AP, 300 Hz to 3 kHz). All stereotactic coordinates were characterized according to their distance to lambda, either in the medio-lateral (ML), dorso-ventral (DV), or antero-posterior (AP) axis. All angles and coordinates were recorded in reference to the azimuthal plane at lambda (Paxinos and Franklin, Nixdorf 2007 stereotaxic). The Neuropixels probe was inserted either tangentially in the superior colliculus (SC) from the back (Figures S1C-F, antero-posterior insertion (API): 15 to 25 deg, 500 to 1200 μm ML, -100 to -500 μm DV) or from the side (Figures S1J-M, medio-lateral insertion (MLI): 20 deg to 30 deg, -100 to -500 μm DV, 0 to 900 μm AP) (Sibille et al., 2021). In the zebra finch, insertion was performed at 40 deg (Figures S7A-D, in reference to lambda: 3000 to 3800 µm ML, -4250 to - 5000 µm DV, 4000 to 4800 µm AP).

### Pupil tracking

To monitor pupil position and dilation in awake recordings, we captured the contralateral eye on a camera (Basler acA 1300) equipped with a zooming lens (850 nm bandpass filter, ThorLabs) using a custom written pupil tracking software. The eye was illuminated with an infrared light source (ThorLabs LZ1-10R602). To avoid interference between the camera with the visual stimulus, eye tracking was performed via a dichroic mirror (Semrock, FF750-SDi02-25×36) that was placed between the eye of the animal and the stimulus screen. The pupil size and position were extracted via DeepLabCut (Mathis et al., 2018) and analyzed using custom-written scripts in Python (Figure S1N/O).

### Visual stimulations

Visual stimuli were generated in Python using the PsychoPy (Peirce, 2008) toolbox. The visual stimuli were displayed on either a calibrated screen (Dell, refresh rate = 120 Hz, mean luminance = 120 cd/m², for the pharmacological recordings, n = 9, Figures 1 and S4, for the awake recordings, n = 3, Figure S1, and half of the recordings in finches) or a calibrated and warped projector image (EPSON projector, refresh rate = 60 Hz, mean luminance = 110 cd/m²). This latter projector image was reflected into a plastic spherical dome screen (EBrilliantAG, IP44, diam = 600 mm) upon a plexiglass reflecting half bowl (Modulor, 0260248). A layer of broad-spectrum reflecting paint was previously applied inside the plastic dome (Twilight-labs). Calibration of the dome warping was done with the Meshmapper software (Paul Bourke). The projector image covered an area of 180 deg * 110 deg which could be re-positioned laterally. We presented the sparse noise visual stimulus on a grid of 36×22 squares with a grid resolution of 5 deg. The sparse noise targets were either dark (on light background) or light (on dark background). We presented three different sparse noise target sizes (5, 10 and 15 deg with respectively [n_target per frames, n_trial per positions] = [6,50], [4,30], [2,20]). Every sparse noise target was displayed for 100 ms. We also presented moving bars (white bars on black background, 10 deg in size, 24 directions, fixed speed of 90 deg/s), chirp stimulus (full-field). The timings of the visual stimuli were marked by stimulus-locked synchronizing signals.

### Pharmacological applications

In the pharmacological experiments, the visual cortex was removed during surgery to avoid any direct visually driven corticotectal activity in the SC. The injector (Drumond, Nanoject II) was inserted vertically in the top part of the SC before recording, to decrease movement-related artifacts (lambda: 500 to 1000 µm ML, 200 to -200 μm AP, 1100 to 1400 μm DV). Approximately 250 nL of different pharmacological cocktails were injected (Figure S5B). Cholera Toxin subunit B, Alexa 488 Conjugate (C22841, Invitrogen) was added to the muscimol solution (Abcam, ab120094, 2.5 mM in PBS). Dextran (Fluorescein, 10,000MW, anionic, D1821) was used in the synaptic blocker mix (Muscimol, NBQX Biozol-HB0443, and D-AP5 Biozol-HB0225 at 2.5 mM, 2.5 mM, 5 mM, respectively, in PBS). Muscimol was injected and allowed to diffuse for 5 minutes before the same stimulus set was repeated. In the successful double-injections pharmacological experiments (n=3/6), first, the injector was removed slowly during the Muscimol period and then the injector’s solution was exchanged with the mixture of synaptic blockers and replaced carefully in a similar location for the second injection (Figure S4). A last repeat of the stimulus presentation followed at the end. Finally, at the end of all pharmacological experiments, a small volume of TTX (∼ 15 μL, Biozol-HB1034, 100 μM in PBS) was applied in the contralateral eye to abolish all remaining visually-driven retinal activity.

### Data analysis

Data analysis was performed in Python (www.anaconda.com) except for Kilosort2 (KS2 (Pachitariu et al., 2016)), a MATLAB (www.mathworks.com) package for spike sorting electrophysiological data, all clusters were manually cured in Phy2. Statistical tests were performed with the Wilcoxon rank-sum test for unpaired samples and with the Wilcoxon signed-rank test for paired samples. Population results are indicated as mean+-standard deviation if not stated otherwise.

### Multi-unit activity extraction

For MUA analysis, the common average reference was applied to the AP band bandpass filtered (Butterworth filter order 2, 0.3 to 3 kHz), where each event is extracted using custom-made Python scripts. Spike detection was performed for each channel independently at a threshold of 4 standard deviations of the AP band (double side detection).

### Spike sorting

Kilosort2 (KS2, https://github.com/MouseLand/Kilosort) was used for spike sorting to produce isolated single-unit clusters followed by manual curation using Phy2 (https://github.com/cortex-lab/phy). Double-counted spikes were removed for each cluster (within ±0.16 ms (Siegle et al., 2021)). Furthermore, between-unit overlapping spikes which produce above chance zero-lag peaks in the cross-correlograms (CCG, peak windows ±0.5 ms) were re-evaluated individually in Phy2 to either refine or drop the problematic cluster(s). Inter-spike-interval (ISI) violations were calculated as the ratio of the spikes within the refractory period (± 1.5 ms) to the total number of spikes. Units with ISI > 0.05% were removed. Furthermore, isolation distance was used as another quality metric to guarantee well isolated clusters (isolation distance > 10 a.u, Figure S3). Other quality metrics, e.g. silhouette score, were calculated using the ecephys spike sorting pipeline (https://github.com/AllenInstitute/ecephys_spike_sorting). We included single unit clusters in the analysis that remained stable over the recording duration.

### Waveform classification

A waveform classification approach was applied to distinguish action potentials from neurons in the vicinity of the Neuropixels probe. For each single-unit cluster, we calculated the multi-channel waveform (MCW) by spike-triggered averaging the raw AP signal on all available spike times (up to 50 000 spike times ±10 ms time window), following an offset correction for each channel. The MCW is therefore the spatiotemporal profile of the AP signals. Afferent axons and somatic signals could be classified based on their distinct waveform (Figure 1D) which allowed us to classify all clusters into afferent axon AP vs somatic AP. We used a two-step approach for this classification. First, a custom-written graphical-user-interface (GUI) was used to manually label the cluster. This GUI was based on (1) the characteristic presence of axonal and dendritic negative peaks within 3 ms (Figure S2), and (2) the possible presence of the axonal path in API (Figures S1E-G). In a second step, we compared our manual classification to an automatized classification using a Gaussian mixture model (GMM) on a principal component projection of classical waveform features (Figure S2A). Using the MCW, the spatial spread of interest (Σ) was estimated by the number of neighboring channels with amplitudes >15% of the cluster maximum at the best channel (BC). This window was interpolated (101 times), smoothed (Gaussian blur 0.1 ms), time-sliced (pre-trough period of 0.6 ms and post-trough period of 3 ms), re-normalized and trough-aligned for more reliable classification. All slope measurements were defined as the 80^th^ percentile values of the observed peaks. All 14 features measures were averaged across the channels of the defined spatial spread (Figure S2B). Additionally, a smaller portion of the obtained axonal clusters from KS2 was detected on their putative dendritic responses (Figure S2D, right). These clusters were discarded as their detection occurred on the postsynaptic dendritic responses of SC neurons and not on the action potential of the retinal axons.

### Detecting afferent axons in Neuropixels datasets

The available default waveform plots generated by KS2 can be used to identify afferent axonal waveforms. To do so, the waveforms plotted in make_fig.m should be sorted based on the value of their second trough at 1.5 ms (from the standard 2.73 ms time window KS2 works with). Consequently, a five minutes long recording would be sufficient to assess whether the given insertion captures afferent axonal waveforms. During curation, the rejection criteria such as “multiple spatial peaks” and “too large spread” (Siegle et al., 2021) should be minimized to increase the chances of identifying axonal waveforms from the dataset.

### Afferent axon conduction velocity

Our method allows the recording of AP signals traveling along the afferent axonal path in the multi-channel waveform and therefore we could estimate the conduction velocity of the AP. Due to the design of the analog-digital conversion of the Neuropixels probe, we had to incorporate an interpolation step before estimating the conduction velocity from the MCW. Briefly, a time correction between adjacent channels was performed to compensate for the serial delay of 2.78 μs between two consecutive channels within each analog-digital converter (personal communication with J. Colonell). Axonal conduction velocity was then estimated on the corrected MCW using a minimal window of 8 channels below the BC (4 channels for zebra finch data) and 10 channels above the tip of the Neuropixels probe. The time of the action potential in the axonal path (−2.5 to -0.1 ms) was detected if its amplitude reached 4*std of the measured baseline (−5 to -2.5 ms). The conduction velocity was estimated by fitting a line through each channel local minima (Figure S1G). This analysis was performed independently for each of the four electrode columns of the Neuropixels probe and the best fit was used as the measure for conduction velocity. Conduction velocity estimates with R^2^ below 0.8 were not included (Figures S1).

### Electrically coupled neighboring retinal ganglion cells

Putative electrically coupled RGCs can be identified based on the presence of characteristic double peaks in the spike train cross-correlograms (Mastronarde, 1983). To identify significant double peaked CCGs, we estimated the baseline between -10 to -5 ms and the peaks on both sides of the zero-lag (−2.5 to -0.5 ms and 0.5 to 2.5 ms). RGC-RGC pairs were considered coupled when both peaks were significantly different (> 3 * std) from baseline (Figures S5B/C).

### Synaptic contact field of afferent axonal arbors

The high-density of recording sites enables to identify the spatial location of the electrical signals of RGC axons on the probe and hence their anatomical location within SC. Importantly, the waveforms of RGC axons also contains the post-synaptic response of SC dendrites (Figures 1D and F) which we used as a proxy for the anatomical location where the RGC axonal arbors make synaptic contacts with SC neurons. Since the Neuropixels probes is organized in four columns of electrodes we could measure the spatial location both along and across the probe (Figures 2A, bottom right) and we define this area on the probe the axonal arbor synaptic contact field (AF). To characterize the spatial position of AF we fitted a two-dimensional Gaussian function to the two-dimensional representation of the synaptic contact field (Figure 2A, bottom-right). This Gaussian fit was necessary because some of the RGC AFs were only partially covered by the recording sites on the probe, e.g. the example in Figure 2A. To fit the AFs we assumed fixed widths of the Gaussian functions and this only optimized the x and y position of the Gaussian by least squares fitting. The AF center position was estimated from the Gaussian fit and could be located close to the electrode border or even outside of the recording sites (Figures 2A and S5/S7).

### Comparing retinal ganglion cell mosaics at the level of dendrites and axons

To compare the spatial organization of the RGC mosaic at the level of receptive fields (RF) within the retina with the organization of the RGC mosaic at the level of the RGC axons within the SC/OT we estimated the RF and AF centers using the center-of-mass measurement (Figure 2C, RF centers = circles, AF centers = crosses). The ensemble of RF and AF center positions were subsequently used to characterize the RGC mosaics with the following two measurements. We calculated the Euclidean distance between the RF centers of RGC pairs (dRF) in the visual space (deg) and the distance between the AF centers (dAF) in the SC space (µm) (Figure 2D). To test for hexagonality of the mosaics we used the Delaunay and Voronoi tessellations (Zhan and Troy, 2000) and estimated the angles of the Delaunay triangles (Figure 2E). For directly comparing the similarity between the RF and AF mosaics we first had to transform the AF mosaic from SC space (µm) into the visual space of the RF mosaic (deg). We achieved this by linearly scaling and rotating the center positions of the AF mosaic such that the summed distances between RF and AF positions were minimized. Important to note, this alignment step does not change the geometric organization of the AF mosaic. From this transformed AF mosaic we calculated the distance between RF and AF centers of individual RGCs. We divided this RF-AF distance by the mosaic spacing factor, which was estimated as the median RF distances between nearest RGC neighbors. Thus, a mosaic spacing of one describes the distance between two neighboring RGCs and a value of zero the situation when the centers are overlapped (Figure 2G). To assess the similarity in the geometrically organization of the RF and AF mosaics we calculated the Euclidean distances between RFs (dRF) and AFs (dAF) of pairs of RGCs (Figure 2H) and estimated the enclosed angle within triangles between RGCs (Figure 2I).

### *In vivo* connectivity analysis

Monosynaptic connections between RGC axons and SC neurons were detected using established methods (Bereshpolova et al., 2020; Denman and Contreras, 2013; Reid and Alonso, 1995; Usrey et al., 1998) on the jitter corrected cross-correlograms (CCGs) (Figure S6A-C) based on statistically significant peaks at synaptic delays (+0.5 to 3.5 ms, purple) above the baseline (Figure S6A, -3.5 to 0 ms, green). Peaks had to extend over the threshold for at least 5 consecutive time bins (0.1 ms resolution). The cross correlations were calculated using the pycorrelate package (https://github.com/tritemio/pycorrelate) (Figure S6). Spike times over the entire recording were used in the CCG analysis. The jitter correction was required to remove stimulus-evoked common input. To estimate the jitter correction, we followed established approaches (Denman and Contreras, 2013; Smith and Kohn, 2008). Briefly, we calculated a jittered version of each spike train by randomizing all spike times within consecutive 10-15 ms windows (Smith and Kohn, 2008). We then calculated the cross-correlation between a pair of neurons both for the original (raw CCG) and the jittered spike train (jittered CCG). Subtracting the jittered CCG from the raw CCG results in a jitter-corrected CCG.

### Efficacy and contribution

Synaptic efficacy and contribution measures of connected pairs were estimated using standard approaches (Bereshpolova et al., 2019; Reid and Alonso, 1995; Swadlow and Gusev, 2002; Usrey et al., 1999). Briefly, efficacy was estimated from the jitter corrected CCGs by dividing the area of the CCG peak (peak window: 0.5 to 3.5 ms, Figure S6A, purple) by the total number of presynaptic spikes. Thus, an efficacy measure of 1 (100%) would reflect that for each presynaptic spike, a postsynaptic spike could be detected. To estimate the contribution, we counted the number of SC spikes that were preceded by a retinal afferent spike, in a time window between -3 to -0.5 ms, and divided this number by the total number of SC spikes. A contribution of 1 (100%) indicates that all spikes of an SC neuron are preceded by retinal activity.

### Receptive fields

Spatiotemporal receptive fields (STRF) were calculated from the peristimulus time histogram (PSTH) for each stimulus position on the 36×22 grid with a 1 ms resolution, resulting in a 3D matrix of visually evoked activity (x-space, y-space, time). The spatial receptive fields were calculated via spike-triggered average (STA) and by using the receptive field at lag -1 as the corresponding onset RF (Kremkow et al., 2016). RFs were interpolated by a factor of two using the 2D-cubic-interpolation function from the SciPy package. RF size was estimated from both responses to light and dark sparse noise stimuli. The RF overlap index was calculated as the number of overlapping pixels in both ON and OFF RFs, defined as 40% of the RFs maximum. The RF distance was calculated by the Euclidean distance between the RF centers. The similarity of the STRFs between pairs of neurons was estimated by the correlation coefficient. For this analysis, the STRFs were calculated with a 100 ms resolution (Figure 5A). Only RFs with high signal-to-noise ratio (SNR >10) were included in the analysis.

### Orientation tuning

To determine orientation tuning, we quantified the responses to moving bars as the maximum response of the PSTH in each exposed direction. The obtained tuning curve was first interpolated on 30 points before fitting with von Mises function (Kremkow et al., 2016) using the least square optimization function from SciPy (Figures 4C-D and Figures S8).

## Data availability

The datasets that support the findings of this study are available from the corresponding author on reasonable request.

## Code availability

All Python code will be made available upon request.

**Figure S1.**
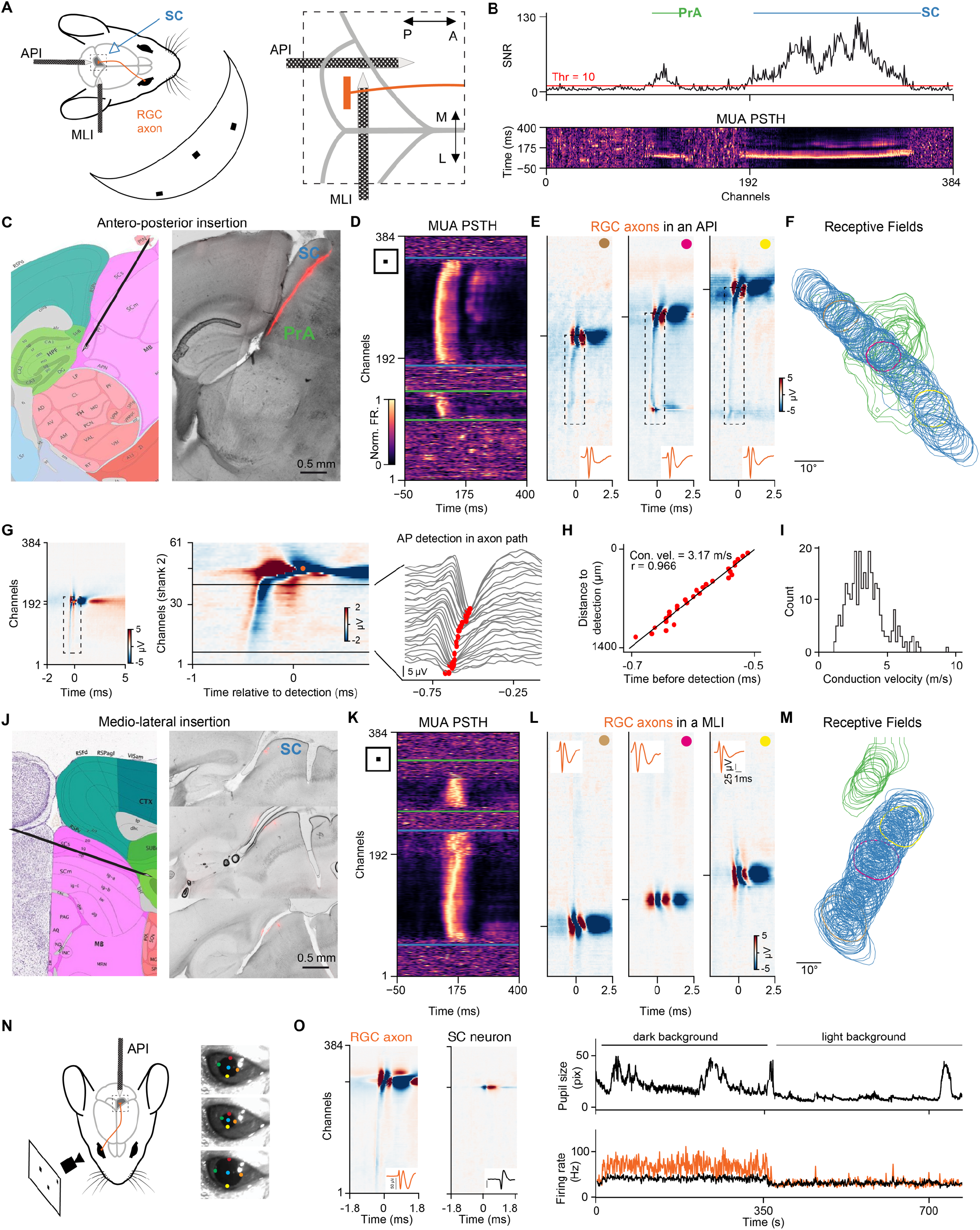
Recording RGC axons and SC neurons in the mouse *in vivo*, related to Figure 1. (A) Schematic of the mouse SC with the axonal projections of retinal ganglion cells (RGC) and visual dome setup (left). Zoom onto the Neuropixels probe implantations along the anterior posterior axis (API) and mediolateral axis (MLI) (right). (B) Visually evoked multi-unit-activity (MUA). Peri-stimulus time histograms (PSTH) to white sparse noise on black background (bottom) and its corresponding signal-to-noise ratio (SNR) (top). In this recording two regions exhibit visually driven activity: SC and pretectal areas (PrA). (C) Sagittal brain slice with a DiI staining of the Neuropixels recording track from an API and the corresponding location in the Allen Institute Common Coordinate Framework (CCF). (D) MUA PSTH to the sparse noise stimulus for the API in C. The SC response (between the blue horizontal lines) covers the upper channels, the lower region is likely within the pretectal areas (green). (E) Multi-channel waveforms (MCW) of three selected RGC axons from the recording shown in D. The dashed black rectangle highlights the axonal action potential (AP) propagating along the axonal path. The best channel (BC), i.e. the channel with the largest waveform amplitude, is indicated by the horizontal tick, and the insets show the RGC axonal waveform at their respective BC in orange. (F) MUA receptive fields (RF) in each visually driven channel: SC (blue) and PrA (green). RFs at the BC of the example single units shown in E are plotted in their respective color. (G) RGC axonal AP conduction velocity estimation in APIs. The AP propagation along the axon path (dashed rectangle) is evident in MCWs of RGC axons recorded in API, left. Close up view on the MCW in a single electrode shank (middle) with the detection of the AP in the axonal path by the local minima of the waveform in each channel (right, red dots). (H) Axonal conduction velocity was estimated from a linear fit to the AP timepoints along the axon path (red dots in G, right). (I) Histogram of RGC axonal conduction velocities across multiple recordings (mean conduction velocity = 3.5±1.3 m/s, n = 283 RGC axons, for RGC axons with R^2^ of linear fit > 0.8). (J-M) Example recording along the mediolateral axis. Same format as for API shown in. C-F. Note that RGC axon paths are not visible in MCWs of RGC recorded in MLI because the Neuropixels probe is not in line with the RGX axons paths in this recording configuration. (N) Recording RGC axons in SC of awake mice. Shown is the API recording configuration used to record from awake SC together with example images of the pupil tracking. The circles on the eye images schematically indicate the markers used for pupil position and size estimation. (O) Example RGC axon and SC neuron recorded in awake SC (left). Pupil size during different stimulus conditions (top) and the neuronal activity (right) of an RGC axon (orange) and an SC neuron (black).

**Figure S2.**
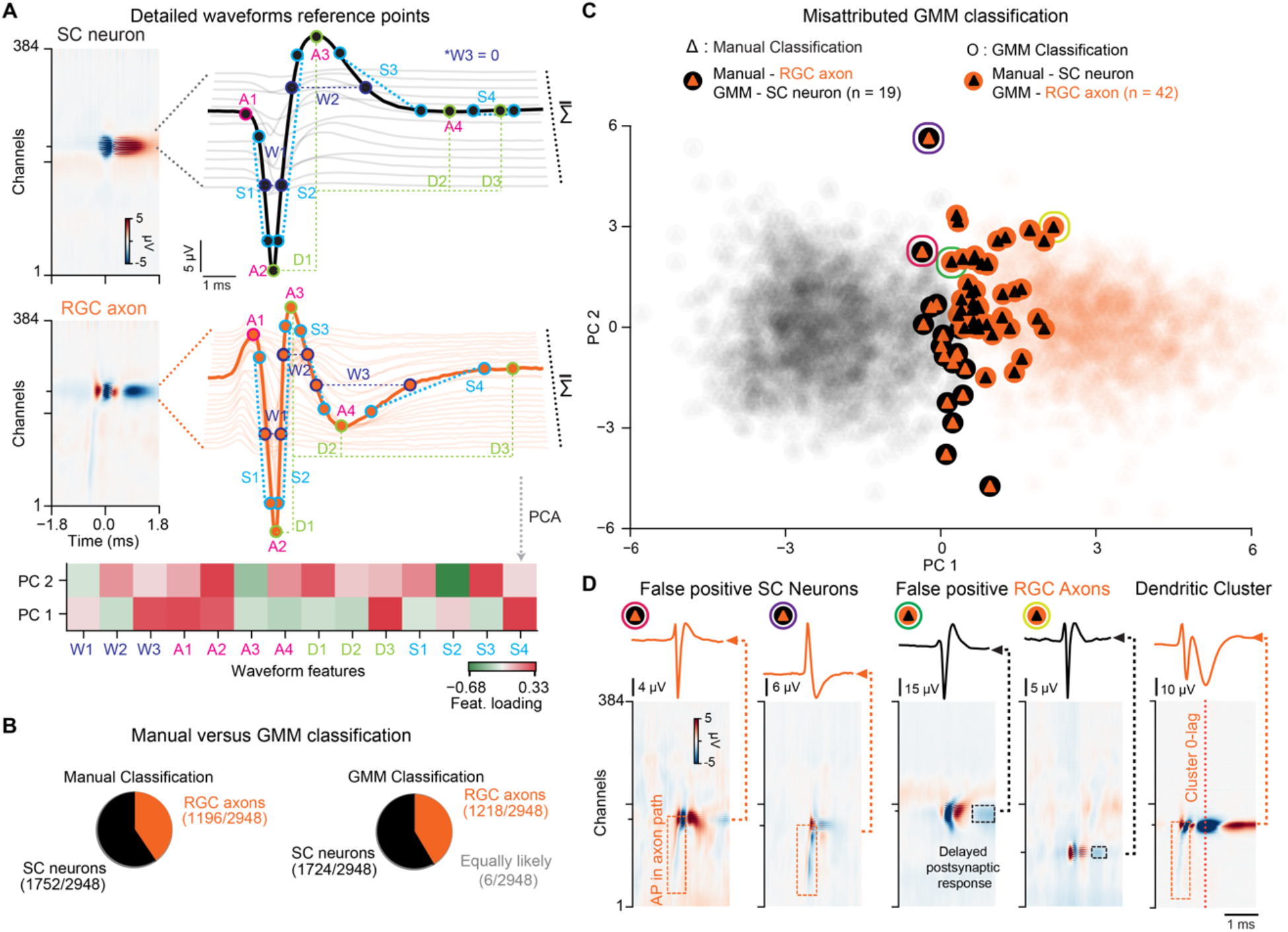
RGC axon and SC neuron waveform classification, related to Figure 1. (A) Example waveforms of a somatic action potential from a single SC neuron (top) and from an RGC axon in SC (middle). The multi-channel waveforms are shown as colormaps together with traces of several single-channel waveforms within the spatial spread of interest. The waveforms were characterized by the following measurements: W1 : half peak width of the detected negative peak (DNP), W2 : half peak width of positive peak after detection (PPAD), W3 : half peak width of second negative peak after detection (SNPAD), A1 : amplitude of the positive peak before detection (PPBD), A2 : DNP’s amplitude, A3 : PPAD’s amplitude, A4 : SNPAD’s amplitude, D1 : difference DNP to PPAD, D2 : difference DNP to SNPAD, D3 : difference DNP to baseline, S1 : depolarization slope, S2 : repolarization slope, S3 : recovery slope PPAD to SNPAD, S4 : recovery slope SNPAD to baseline. The loading of these different features on the two first principal components (PCs) is represented in a heatmap (bottom). The two first PCs were used in the Gaussian mixture model (GMM) for classification. (B) Pie charts showing the results of the GMM vs. the manual classification. (C) Scatter plot of PC1 and PC2 projections (top) of the manual classification (blurred). Misclassified GMM clusters are shown as large colored disks. Four examples (purple, red, green, yellow circle on top, corresponding to the examples in d) illustrate misclassified clusters and their corresponding MCW. (D) False positive SC neurons are identified by the presence of an axonal tail (orange dashed squares) in the MCW (two left). False positive RGC axons are identified by the presence of an unexpected negative peak after detection (black dashed squares two examples in the middle right). Waveforms that were detected on the postsynaptic response of the RGC axon waveform were identified as dendritic clusters and excluded from the dataset (right).

**Figure S3.**
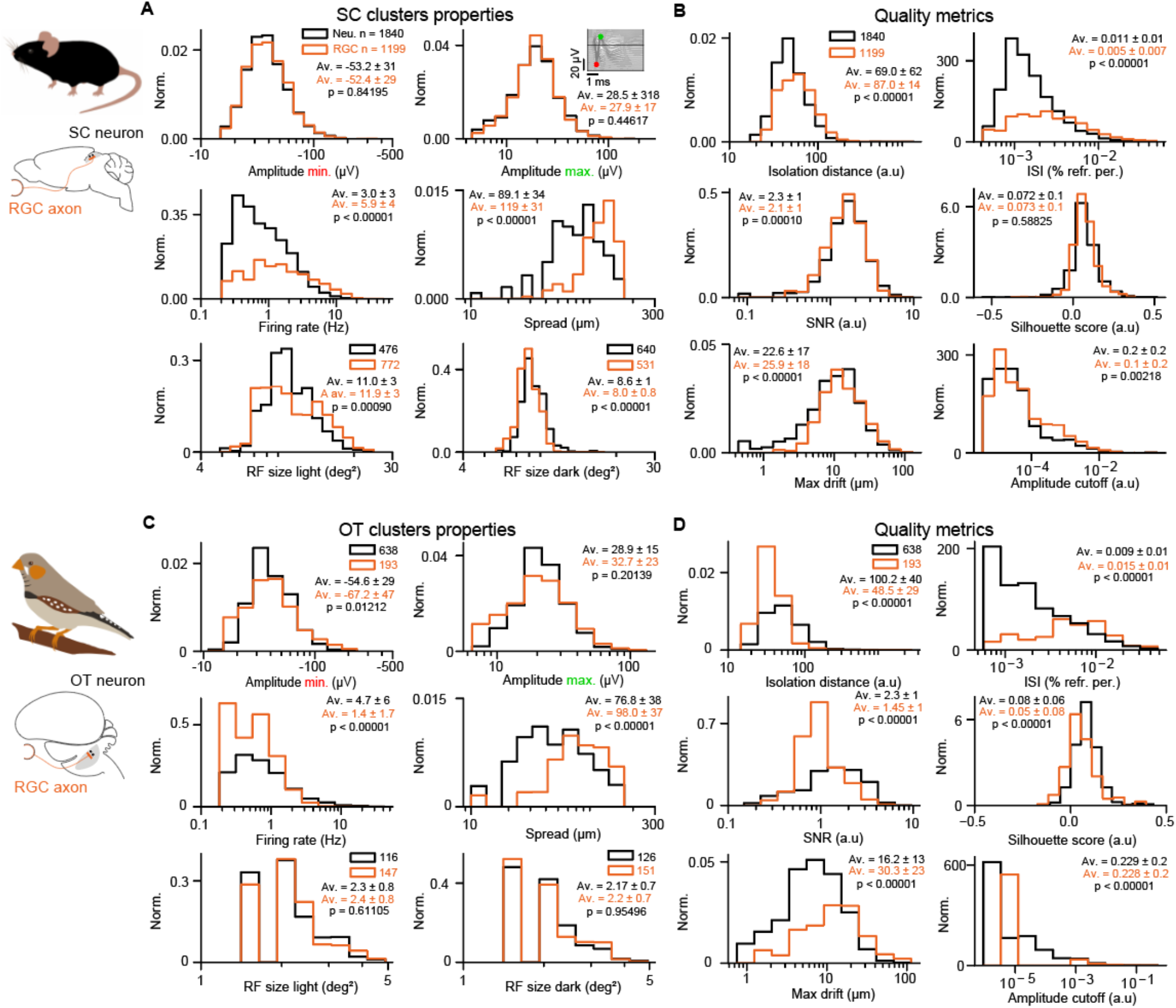
Properties and quality metrics of single units in the mouse superior colliculus and zebra finch optic tectum, related to Figures 1 and 6. (A-B) Overview of the mouse dataset. (C-D) Overview of the zebra finch dataset. (A) Cluster properties in the mouse: Negative (top left) and positive (top right) amplitudes of the waveform are estimated from the MCW, all shown on log-normal scale. Firing rate (FR) (middle left) and waveform spatial spread (middle right) in both populations of RGC axons and SC neurons show significant differences: RGC axons exhibit a higher FR and a larger waveform spread. Receptive field (RF) sizes, obtained from only clusters with high signal-to-noise RFs (SNR > 10) were estimated using both light (left) and dark (right) sparse noise. (B) Single unit quality metrics. RGC axons and SC neurons have similar quality measures. (C) Cluster properties in the zebra finch dataset, same format as in A. (D) Quality metrics in the finch dataset, shown as in B.

**Figure S4.**
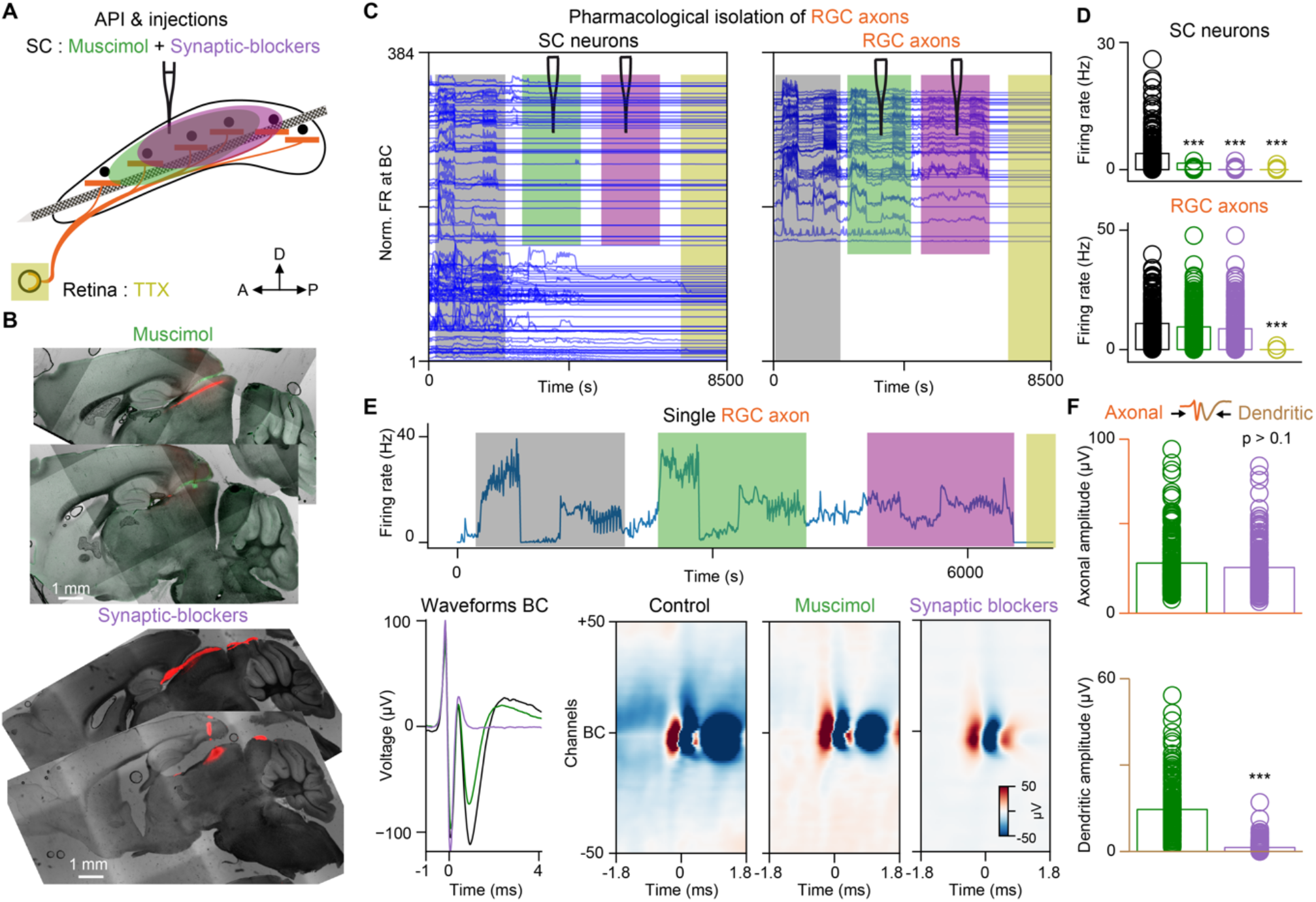
Pharmacological confirmation of RGC axons in the SC, related to Figure 1. (A) Schematic showing the pharmacological injections: Muscimol (green), synaptic blocker (purple) and Tetrodotoxin (TTX, yellow). (B) Sagittal brain slices illustrating the injections of Muscimol (left, green) and synaptic blocker (right, red) with DiI staining from API insertions (top, red). (C) SC neuron activity, aligned to the BC. RGC axons recorded in the same region aligned to their BC (right). The pharmacological injections are indicated by different colors. Activity of SC neurons decreases following Muscimol injection, while the activity of RGC axons remains stable. (D) Quantification of firing rate changes of SC neurons and RGC axons during the different pharmacological conditions. Control (black): SC neurons = 3.8±4.2 Hz, RGC axons = 7.9±10.6 Hz; Muscimol (green): SC neurons = 1.5±2.6 Hz, RGC axons = 7.2±9.3 Hz; synaptic blocker (purple): SC neurons = 0.03±0.16 Hz, RGC axons = 8.67±8.5 Hz; TTX (yellow): SC neurons = 0.01±0.13 Hz, RGC axons = 0.1±0.006 Hz. n = 224 SC neurons, n = 215 RGC axons. (D) A single RGC axons during the entire recording (top) illustrating the stability of RGC axon detection even after application of the synaptic blockers. The waveform of the same RGC axons is shown at its BC (bottom left) and across 100 channels (bottom right) for the control (black), Muscimol (green) and synaptic blockade (purple) condition. Note that the postsynaptic dendritic rebound (second trough in the RGC axon waveform) is abolished after the application of synaptic blockers. (E) Quantification of the axonal (top) and dendritic signal amplitudes (bottom) during the application of Muscimol (green) and synaptic blockers (purple) (axonal amplitude: Muscimol = 28.8±14.5 µV, synaptic blocker = 26.3±13 µV, p > 0.1; dendritic amplitude: Muscimol = 14.5±9.04 µV, synaptic blocker = 1.39±1.9 µV; n = 203 RGC axons). *** = p < 0.001.

**Figure S5.**
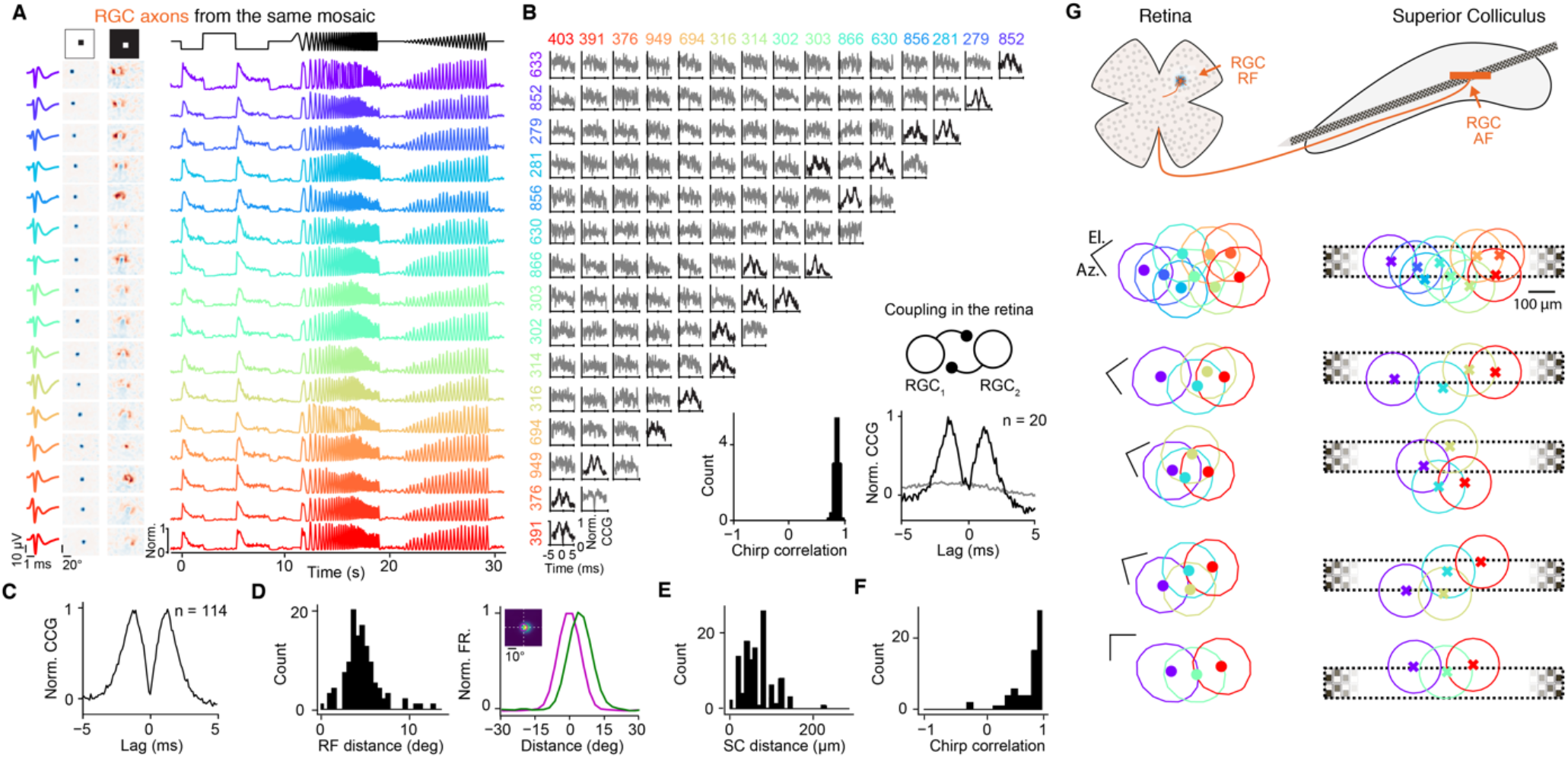
*In vivo* recording of retinal axon mosaics, related to Figures 1 and 2. (A) Simultaneously recorded RGC axons in mouse SC from the same functional retinal mosaic (OFF-transient). Shown are: the waveforms at the best channel (left column), RFs for both dark (middle left column) and light (middle right column) sparse noise targets and the responses to the chirp stimulus (right). Note the classical OFF-center, ON-surround RF organization and the OFF-transient responses during the chirp stimulus for all RGCs. (B) Cross-correlograms (CCGs) between the RGC axons shown in A. Note: neighboring RGCs can show double peaks in the CCGs (black traces) that are characteristic for coupling between RGCs in the retina (Mastronarde, 1989). The insets (bottom, right) show the histogram of the correlation coefficient of the chirp responses between pairs of putative coupled RGCs from this mosaic (mean correlation = 0.93±0.02, n = 20 coupled pairs) and the population average CCG of RGC pairs with significant coupling (black, n = 20 coupled pairs) and uncoupled pairs (gray). (C) Average CCG of coupled RCGs from multiple experiments (n = 114 RGC pairs, n = 32 mice). (D) RF distance of coupled RGC pairs (left) and average RF profiles (right). RGC pairs (green and magenta lines) have partially overlapping RFs (mean RF distance = 4.70±1.95 °, n = 114 RGC pairs). (E) Putative coupled RGCs project to neighboring locations in SC (left, mean SC distance = 65.54±34.53 µm, n = 114 coupled pairs) and the majority show similar functional responses to a chirp stimulus (right, mean correlation = 0.79±0.22, n = 114 coupled pairs). (G) Multiple examples of retina mosaics at the level of RGC dendrites (RF) and RGC axons (AF) measured in mouse SC (n = 4 mice). Scale bars for the RFs: 10 deg.

**Figure S6.**
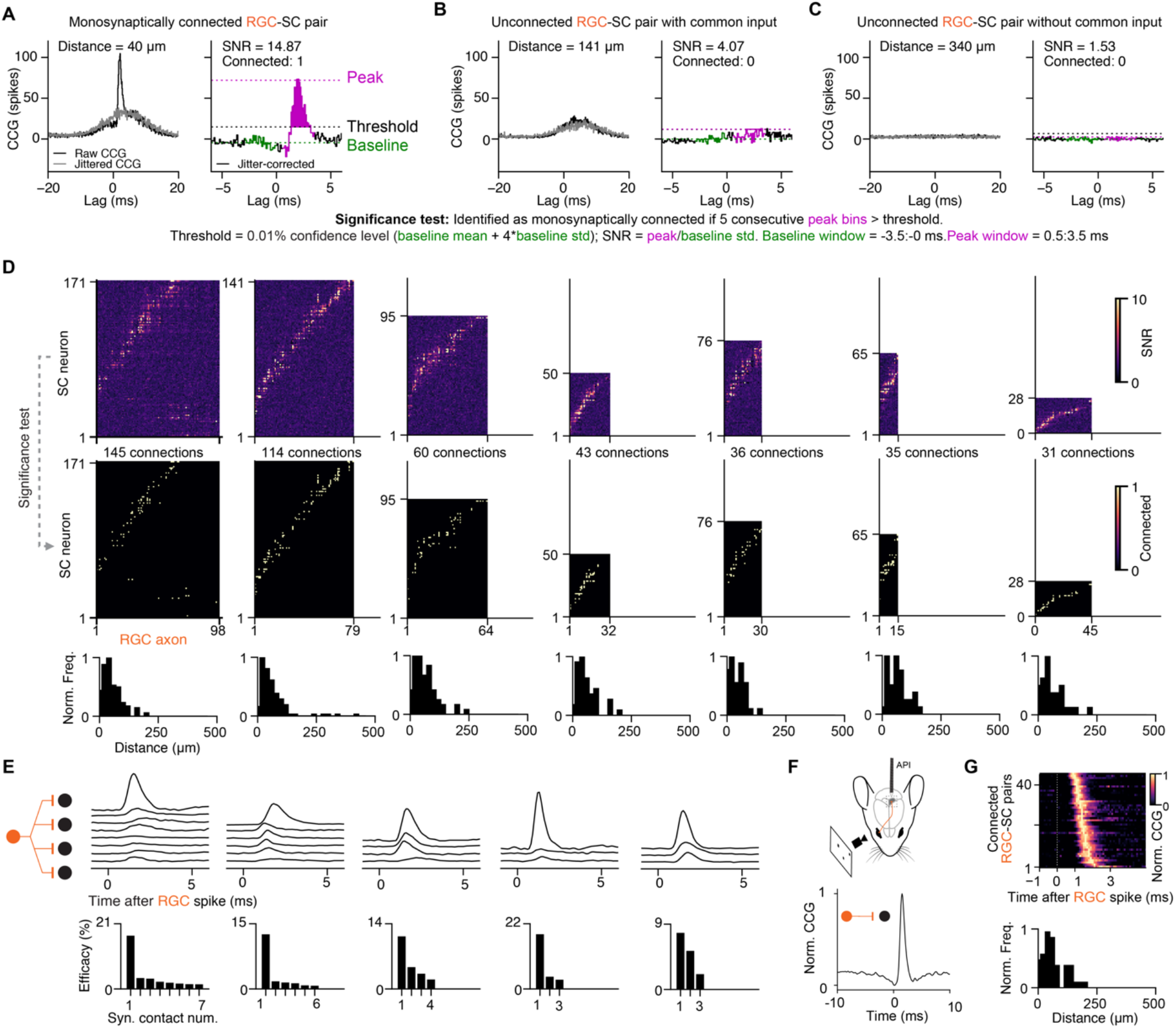
*In vivo* connectivity analysis, related to Figure 3. (A) Example CCG of a monosynaptically connected RGC-SC pair. Left, the black trace shows the raw CCG and the gray trace the jittered CCG. The transient peak at short latency is characteristic for monosynaptically connected neurons while the broader peak is common input. In the jitter-corrected CCG (raw-jitter) the common input is substracted allowing to identify monosynaptically connected pairs (right) by estimating the statistical significance of the transient peak (magenta, see Methods). (B-C) Examples of unconnected RGC-SC pairs, with common input (B) and without common input (C). (D) Connectivity matrices of multiple examples recordings: the CCG peak SNR (top) and statistical identified connections (middle) are shown. The histogram of the distances between connected RGC-SC pairs on the probe is shown below. (E) Example CCGs of divergent connections from single RGC axons to multiple SC neurons (top). The efficacy is non-uniform with few strong and several weak connections (bottom). (F) Example of a monosynaptically connected RGC-SC pair in awake mouse. (G) CCGs of connected pairs across multiple experiments in awake mice, sorted by peak latency (top). Distance between connected RGC-SC pairs on the probe (bottom).

**Figure S7.**
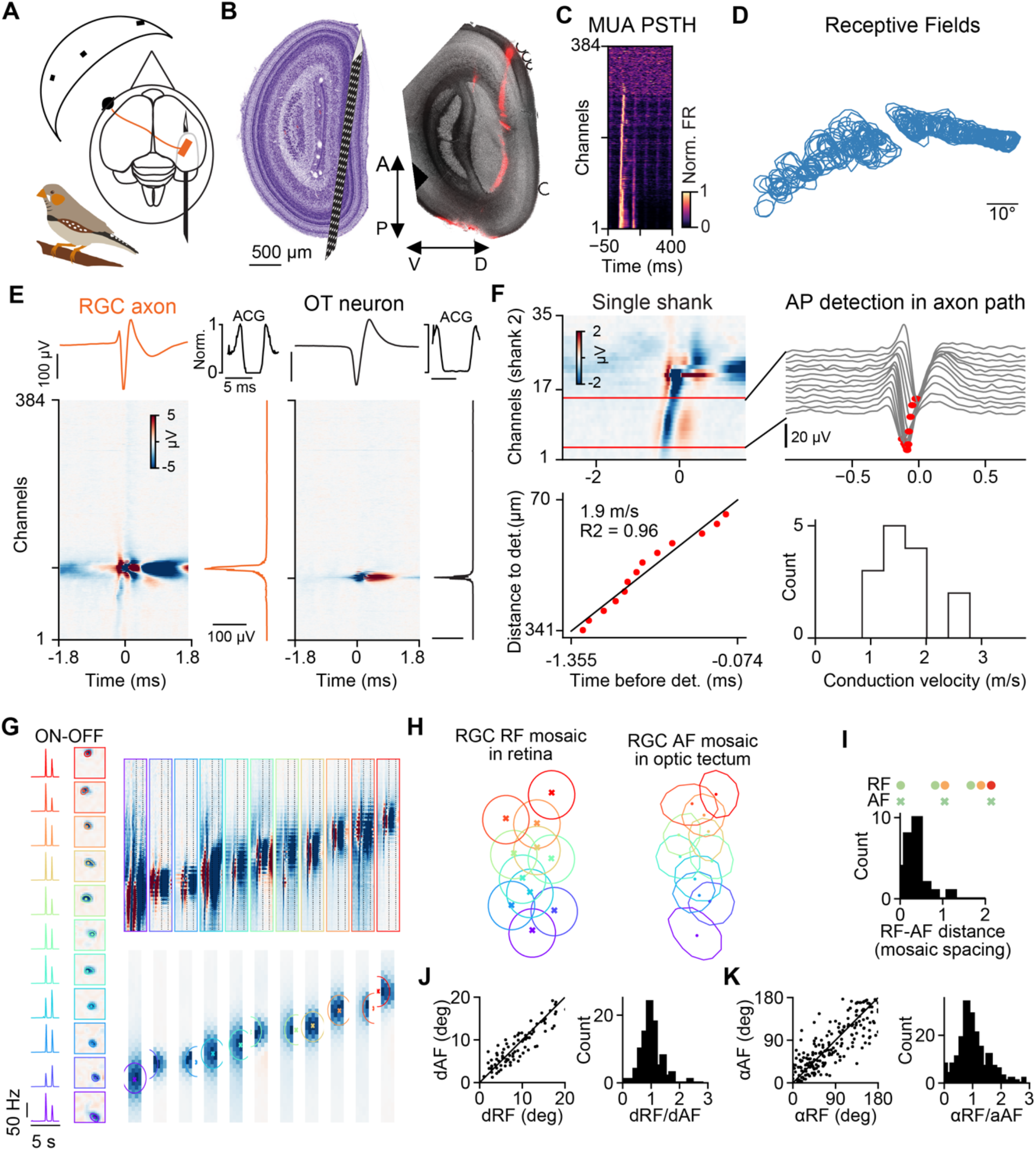
RGC axon mosaics in the optic tectum of zebra finch, related to Figure 6. (A) Retinotectal projections in the zebra finch. (B) Sagittal slices of the OT with DiI staining from an anteroposterior insertion (right), with its corresponding location in the ZEBrA histological atlas (left). (C) Visually-evoked MUA identifies the recording sites located within OT. The colormap shows the PSTH of each channel in response to a sparse noise stimulus. (D) Receptive fields of the MUA shown in C. (E) MCW of an RGC axon and an OT neuron and the corresponding amplitude profile along the probe (right), its waveform shape at the BC (top) with their spike train auto-correlogram (ACG, top right). (F) Closeup view on an RGC axon (top left) in a single electrode shank, with the area of interest (top right) showing the detection of the AP in the axon path by the local minimal (red dots). Axonal conduction velocity (bottom left) was estimated from a linear fit to the AP time points along the axonal path. The fit was performed for each of the four shanks of the probe separately and the best fit was included when R^2^ > 0.75. Conduction velocities across multiple recordings and RGC axons (bottom right, mean conduction velocity = 1.5±0.5 m/s, n = 14 RGC axons, n = 2 zebra finches). (G) Simultaneous measurement of RGCs belonging to the same functional retinal mosaic. RGC functional type was identified using a chirp stimulus (left) and receptive field polarity (left). In this example RGCs were from the ON-OFF type. The corresponding receptive fields (RF, middle) and axonal synaptic contact fields (AF, right) cover a large extend of the visual field and SC tissue. (H) The RFs and AFs mosaic of the example shown in G with their respective contours and colors. (I) Histogram of the distances between RF and AF centers in the unit of mosaic spacing (median RF-AF distance = 0.42±0.25 mosaic spacings, n = 26 RGCs). (J) Distances between RF centers (dRF) and AF centers (dAF) are similar. dRF plotted against dAF and histogram of the dRF/dAF ratios (median dRF/dAF = 1.05±0.33, n = 92 RGC pairs). (K) Angles between RGC triples in the RF and AF mosaic are similar. αRF plotted against αAF and histogram of the αRF/αAF ratios (median αRF/αAF = 1.01±0.85, n = 199 angles).

**Figure S8.**
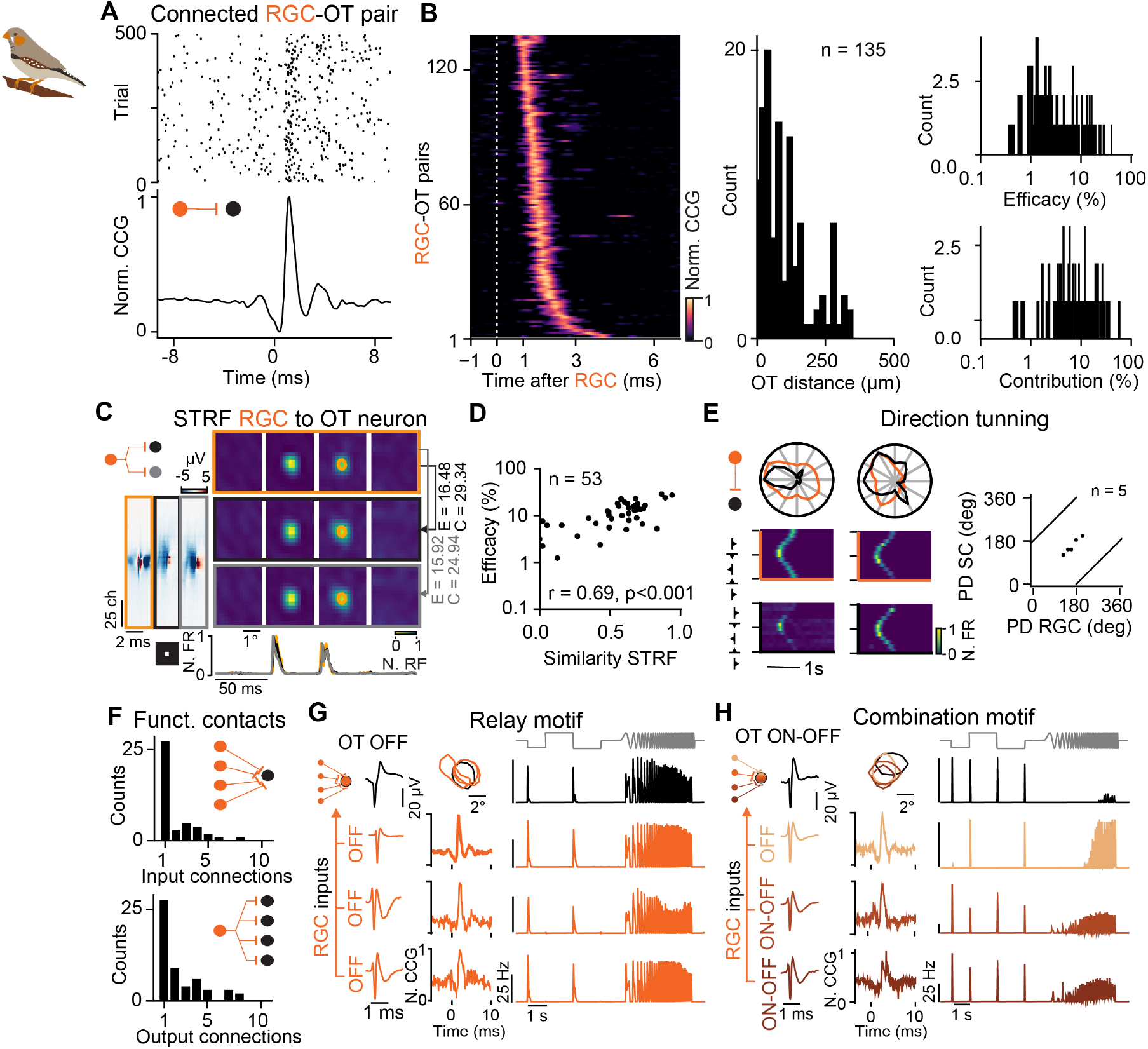
Functional connectivity between retina and optic tectum in zebra finches, related to Figure 6. (A) Example of an identified connection between an RGC axon RGC and an OT neuron. Shown are the raster plot (top) and the corresponding spike train CCG (bottom). (B)Representation of all connected pairs recorded in the zebra finch dataset (left) and the corresponding distance of connected pairs on the probe (middle). The synaptic efficacy (top right) and contribution (bottom right) exhibit log-normal distribution (n = 7 zebra finches, n = 628 OT neurons, n = 193 RGC axons, n = 135 connected RGC-OT pairs). (C) Similarity between visual response properties of an RGC axon (orange) and two postsynaptic OT neurons (black and gray). Shown are the STRF and the corresponding PSTH response (bottom). (D) Correlation between the similarity of STRFs and connection efficacy. (E) Direction tuning of connected RGC-OT pairs. The preferred direction (PD) of RGC axons and their connected postsynaptic OT neurons match (PD difference = 17.26±7.04°, n = 5 connected pairs). (F) Number of measured divergent and convergent RGC-OT connections. (G) Relay motif example, three RGC axons (orange) converge onto one OT neuron (black), all exhibiting similar OFF responses and have overlapping RFs, different waveforms and single-sided CCG peak. (H) Same as in g but for a combination motif example. Here, different RGC response types converge onto an ON-OFF OT neuron.

## Notes

### Competing Interest Statement

The authors have declared no competing interest.

